# Genetic and Epigenetic Features of Promoters with Ubiquitous Chromatin Accessibility Support Ubiquitous Transcription of Cell-essential Genes

**DOI:** 10.1101/2020.11.02.364869

**Authors:** Kaili Fan, Jill E. Moore, Xiao-ou Zhang, Zhiping Weng

## Abstract

Gene expression is controlled by regulatory elements with accessible chromatin. Although the majority of regulatory elements are cell type-specific, being in the open chromatin state in only one or a few cell types, approximately 16,000 regions in the human genome and 13,000 regions in the mouse genome are in the open chromatin state in nearly all of the 517 human and 94 mouse cell and tissue types assayed by the ENCODE consortium, respectively. We performed a systematic analysis on the subset of 9,000 human and 8,000 mouse ubiquitously (ubi) open chromatin regions that were also classified as candidate cis-regulatory elements (cCREs) with promoter-like signatures (PLSs) by the ENCODE consortium, which we refer to as ubi-PLSs. We found that these ubi-PLSs had higher levels of CG dinucleotides and corresponded to the genes with ubiquitously high levels of transcriptional activities. Furthermore, the transcription start sites of a vast majority of cell-essential genes are located in ubi-PLSs. ubi-PLSs are enriched in the motifs of ubiquitously expressed transcription factors and preferentially bound by transcriptional cofactors that regulate ubiquitously expressed genes. Finally, ubi-PLSs are highly conserved between human and mouse at the synteny level, but not as conserved at the sequence level, with a high turnover of transcription factor motif sites. Thus, there is a distinct set of roughly 9,000 promoters in the mammalian genome that are actively maintained in the open chromatin state in nearly all cell types to ensure the transcriptional program of cell-essential genes.

## Introduction

Cells in a multicellular organism share the same genome but interpret it differently to carry out cell type-specific transcriptional programs. Cell type specificity partly resides in maps of chromatin accessibility [1,2], DNA methylation [3], and histone modifications [4], and partly in the levels of regulatory proteins such as transcription factors [5]. Among the three types of maps, chromatin accessibility is a powerful indicator of whether a genomic region may have regulatory functions, while DNA methylation and histone modifications suggest the type of regulatory functions (e.g., promoters, enhancers, or insulators). DNase-seq [6] and ATAC-seq [7] are two widely used techniques for mapping chromatin accessibility, and they have revealed that chromatin accessibility maps are highly variable across cell and tissue types [8–10].

In an early study, we used a DNase-microarray assay to map chromatin accessibility in 1% of the human genome in six cell lines [11] and observed that 22% of the DNase hypersensitive sites (DHSs) were shared among all six cell lines. We called these ubiquitous DHSs and found them to be enriched in promoters and insulators, while the cell type-specific DHSs were enriched for enhancers [11]. It was unclear whether this set of ubiquitous DHSs would remain as a distinct group when chromatin accessibility maps become available for a large number of biosamples, and if so, whether we could discern more biological features for the group beyond the enrichment in promoters and insulators.

As part of the ENCODE Project, we identified a representative set of 2.16 million DNase hypersensitive sites (rDHSs) using the ENCODE data from the ~70 million DHSs identified in more than 500 DNase-seq experiments across diverse cell and tissue types [12]. We further defined a subset of rDHSs with high ChIP-seq signals (defined as Z-score > 1.64) of two key histone modifications (H3K4me3 and H3K27ac) or the chromatin-structure protein CTCF as candidate cis-regulatory elements (cCREs) [12]. cCREs were further classified into groups according to whether they had high signals in these four assays (DNase-seq and ChIP-seq of H3K4me3, H3K27ac, and CTCF) in a particular biosample (i.e., unique tissue sample or cell type) and based on their distances from GENCODE-annotated transcription start sites (TSSs)—cCREs with promoter-like signatures (PLSs), enhancer-like signatures (ELSs) (further divided into TSS-proximal pELS and TSS-distal dELS using a cutoff of 2 kb), and three other groups [12]. When we examined the cell type specificity of cCREs in the 25 human biosamples with data from all four assays, we observed that cCRE-PLSs tended to be active (defined as having high DNase or ChIP-seq signals) in multiple biosamples while cCRE-ELSs tended to be active in only one or a few biosamples [12], consistent with earlier results on promoters and enhancers [8,11,13].

We took advantage of the extensive collection of ENCODE data to revisit our earlier question about ubiquitous DHSs. We defined 15,989 ubiquitous human rDHSs (ubi-rDHSs) and 13,247 ubiquitous mouse rDHSs with high DNase signals in at least 95% of 517 human DNase-seq experiments and 94 mouse DNase-seq experiments, respectively. Confirming our earlier results [11], we found that nearly 60% of ubi-rDHSs were promoters (referred to as ubi-PLSs), and roughly 20% additional ubi-rDHSs were TSS-proximal enhancers (within 2 kb of an annotated TSS). We also found that ubi-PLSs are a set of regulatory elements with distinct properties. Compared with the remaining cCRE-PLSs (called non-ubi-PLSs), ubi-PLSs are highly enriched in CG dinucleotides, a sequence feature that inherently leads to low chromatin accessibility, as revealed by in vitro DNase-seq data [14]. However, ubi-PLSs are also enriched in the motifs of ubiquitously expressed transcription factors and preferentially bound by transcriptional cofactors that regulate ubiquitously expressed genes; thus, the binding of the transcriptional machinery and regulatory proteins is the driving force behind the ubiquitous chromatin accessibility of ubi-PLSs.

Furthermore, ubi-PLSs are highly conserved between human and mouse, and they correspond to the TSSs of cell-essential genes. Thus, there is a distinct set of roughly 9,000 promoters in the mammalian genome actively maintained in the open chromatin state in nearly all cell types to ensure the transcriptional program of cell-essential genes.

## Results

### Three-quarters of ubiquitously DNase-accessible sites are proximal to TSSs

The 2,157,387 human rDHSs previously defined by the ENCODE consortium [12] showed a bimodal distribution in terms of the number of biosamples in which they had high DNase-seq signals (**Figure 1A**). Most of these rDHSs had high DNase signals in one or a few biosamples, while a small but distinct subset of rDHSs had high DNase-seq signals in almost all of the 517 human biosamples included in our study (**Figure 1A**). We defined the 15,989 rDHSs with high DNase signals in 500 or more biosamples as ubi-rDHSs (**Supplementary Table 2A**).

**Figure 1.**
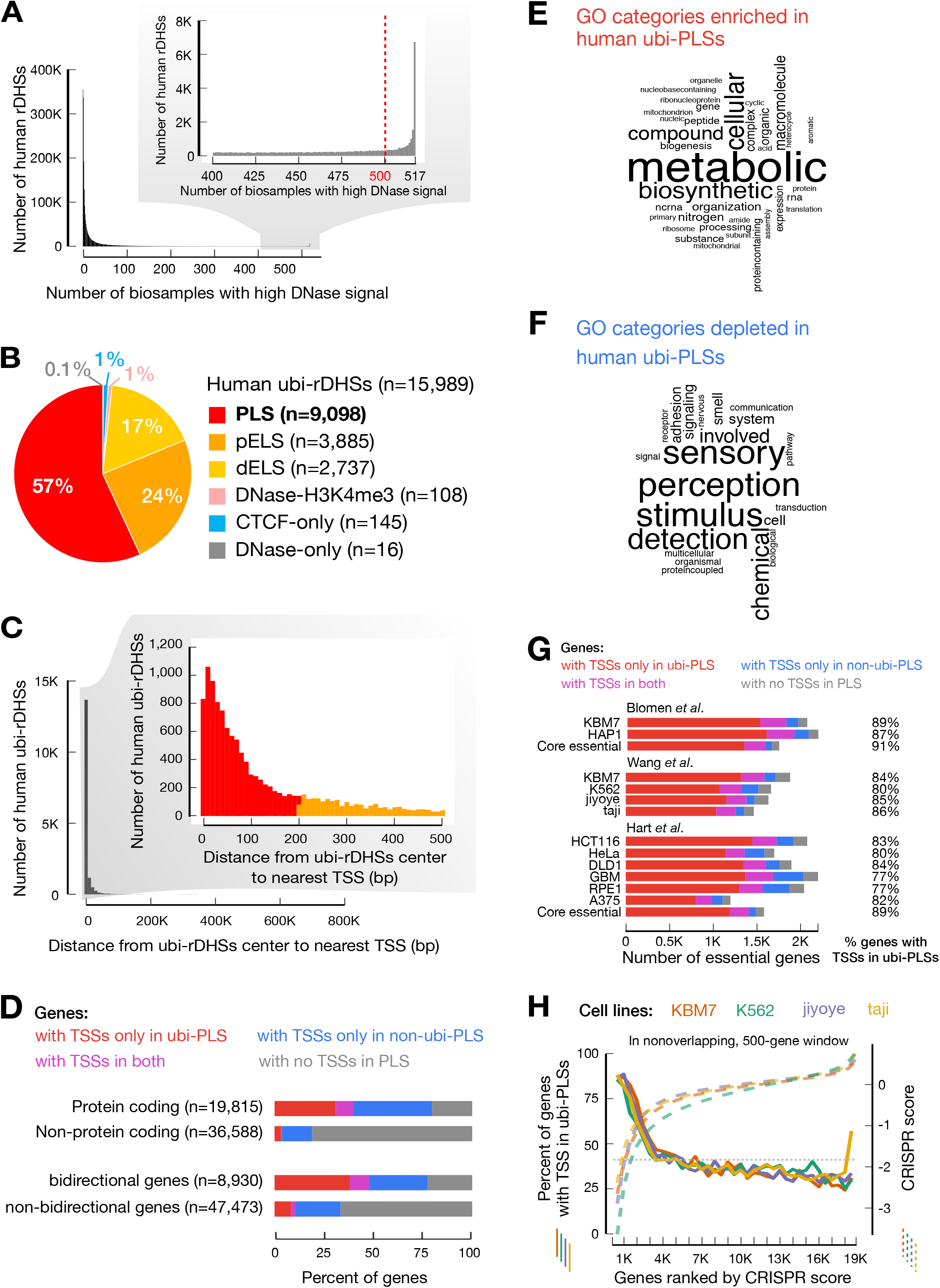
The majority of ubiquitously DNase-accessible regions are active promoters defining essential genes. **A.** Definition of ubi-rDHSs. The histogram shows the number of human rDHSs that have high DNase signals in different numbers of biosamples. rDHSs that have high DNase signals in 500 or more human biosamples (out of a total of 517 biosamples) are defined as ubi-rDHSs. **B**. The majority of human ubi-rDHSs have promoter-like signatures or are TSS-proximal with enhancer-like signatures. The pie chart shows the category of ubi-rDHSs: PLS, rDHSs with promoter-like signature; pELS, TSS-proximal enhancer-like signature; dELS, TSS-distal enhancer-like signature; DNase-H3K4me3, TSS-distal rDHSs with high DNase and high H3K4me3 signals but low H3K27ac signals; CTCF-only, rDHSs with high DNase and high CTCF signals but low H3K4me3 and H3K27ac signals; DNase-only, rDHSs with high DNase signals but low H3K4me3, H3K27ac, and CTCF signals. **C.** The majority of ubi-rDHSs are near a TSS. The histogram shows the number of ubi-rDHSs at a certain distance from the nearest GENCODE-annotated TSS. **D.** Higher percentages of protein-coding genes and bidirectional genes have TSSs overlapping ubi-PLSs than other types of genes. Genes with TSSs only overlapping ubi-PLSs are shown in red; genes with TSSs only overlapping non-ubi-PLSs are in blue; genes with TSSs overlapping both ubi-PLSs and non-ubi-PLSs are in purple; and genes with no TSSs overlapping PLSs are in gray. **E.** Word cloud of the most enriched GO Biological Process terms for genes with TSSs overlapping ubi-PLSs. **F.** Word cloud of the most depleted GO Biological Process terms for genes with TSSs overlapping ubi-PLSs. **G.** A high percentage of cell-essential genes have TSSs overlapping ubi-PLSs. Three groups of bar plots represent cell-essential genes from three different studies, while core essential genes overlap essential genes between different cell lines defined in the studies. Classification of genes in this figure are the same as in Figure 1D. **H.** The TSSs of most top-ranked cell-essential genes are located in ubi-PLSs. Four solid lines show the percentage of cell-essential genes with TSSs overlapping ubi-PLSs, plotted for nonoverlapping, 500-gene windows in four cell lines (y-axis on the left) as functions of the ranks of the genes according to their CRISPR scores. Cell-essential genes are from Wang et al. CRISPR scores (y-axis on the right, dashed lines) are also shown as functions of the ranks of these scores. The horizontal dotted line shows the percentage of all tested genes with TSSs overlapping ubi-PLSs.

ENCODE deemed 926,535 rDHSs—those with high signals of two histone modifications (H3K4me3 and H3K27ac, characteristic of promoters and enhancers, respectively [15,16]) or the chromatin-structure protein CTCF—cCREs, and further classified these cCREs into five groups [12]. Nearly all ubi-rDHSs were cCREs (**Figure 1B**), and 57% of ubi-rDHSs (n = 9,098) were cCREs with a promoter-like signature (PLS; high H3K4me3 and having a TSS within 200 bp of the rDHS center [12]). Additionally, 24% of ubi-rDHSs (n = 3,885) were TSS-proximal cCREs with an enhancer-like signature (pELS; high H3K27ac and having a TSS within 200 bp of the rDHS center [12]). In aggregate, most ubi-rDHSs were within 500 bp of a TSS (**Figure 1C**). We focused our subsequent analyses on the 9,098 ubi-rDHSs that were also cCRE-PLSs, which we denote ubi-PLSs. For comparison, we refer to the remaining 25,705 cCRE-PLSs that were not ubi-rDHSs as non-ubi-PLSs.

### ubi-PLSs define essential genes

The 9,098 ubi-PLSs overlapped 18,187 GENCODE V24 annotated TSSs, which belonged to 9,214 genes (**Supplementary Table 2B, C**). Among these, 7,340 genes have TSSs exclusively overlapping ubi-PLSs, and the other 1,874 genes additionally have one or more TSSs overlapping non-ubi-PLSs. Meanwhile, 13,544 genes have TSSs only overlapping non-ubi-PLSs. The number of genes with both TSSs overlapping ubi-PLSs and other TSSs overlapping non-ubi-PLSs (1,874) is substantially lower than expected (6,732 ± 44, *p*-value < 1 × 10^-4^) if the TSSs were to be distributed randomly while maintaining the number of TSSs in each gene (see **Methods**). Thus, a distinct set of genes use ubi-PLSs to drive their transcription.

Furthermore, ubi-PLSs are enriched for the TSSs of protein-coding genes and bidirectional genes. The vast majority of ubi-PLSs contain the TSSs of protein-coding genes—7,873 (85%) of the 9,214 genes with TSSs overlapping ubi-PLSs are protein-coding genes (defined by GENCODE v24, see **Methods**). Reciprocally, 40% of protein-coding genes have TSSs overlapping ubi-PLSs, while only 4% of non-protein-coding genes have TSSs overlapping ubi-PLSs (Fisher’s exact test, *p*-value < 2.2 × 10^-16^; **Figure 1D**). Among the 8,930 bidirectional genes (defined in **Methods**), 4,265 (48%) have TSSs overlapping ubi-PLSs (**Figure 1E**). Reciprocally, 46% of the 9,214 genes with TSSs overlapping ubi-PLSs are bidirectional (Fisher’s exact test, *p*-value < 2.2 × 10^-16^).

Gene ontology analysis revealed that the 9,214 genes with TSSs overlapping ubi-PLSs were enriched in universal biological processes such as various metabolic, biosynthetic, biogenesis, and translational processes, but were depleted in specialized biological processes such as signaling, communication, sensory perception, response to stimuli, and adaptive immune response (**Figure 1F,G, Supplementary Table 3**). Thus, ubi-PLSs correspond to the promoters of housekeeping genes that perform day-to-day cellular functions.

Cell-essential genes in the human genome were recently defined in several CRISPR screens [17–19]. One study tested 18,166 genes in four cell lines, ranked these genes by their CRISPR scores in each cell line, and deemed the top 10% of the ranked genes essential [17]. We found that 80-86% of these essential genes have TSSs overlapping ubi-PLSs (**Figure 1G**). Furthermore, when we ranked genes by essentiality (CRISPR scores in the range of −5.8 to 2.1 across the four cell lines; the more negative a score, the more essential the gene), we found that the percentages of genes overlapping ubi-PLSs decreased with the rank (**Figure 1H**). Two genes with TSSs overlapping ubi-PLSs, *RPL23A* (Ribosomal Protein L23a) and *CDC16* (cell division cycle 16), were deemed essential in all four tested cell lines (CRISPR scores −3.4 to −5.1 for *RPL23A* and −2.6 to −4.9 for *CDC16).* RPL23A is a component of the ribosome, responsible for protein synthesis. CDC16 is a protein ubiquitin ligase in the APC complex, which governs exit from mitosis via targeting cycle proteins for degradation by the 26S proteasome. Abnormal expression of *CDC16* can lead to diseases such as deafness and early infantile epileptic encephalopathy [20–22]. We also tested the cell-essential genes defined in two other studies [18,19]; 77-89% of essential genes defined in individual cell lines and 89-90% core essential genes (essential in multiple cell lines defined by each study) have TSSs overlapping ubi-PLSs (**Figure 1G, Supplementary Figure 1**). In summary, ubi-PLSs regulate the transcription of genes that maintain essential cellular functions.

### ubi-PLSs are enriched in CG dinucleotides and their epigenetic environments are highly conducive to active transcription

Many promoter sequences are enriched in CG dinucleotides, and high-CG promoters tend to be constitutively expressed across cell types [23,24]. The vast majority of ubi-PLSs have high CG content [24] (normalized CG dinucleotide content ≥ 0.5), while the non-ubi-PLSs show a bimodal distribution (**Figure 2A**). Accordingly, 89% of ubi-PLSs (8,131 of 9,098) overlap CpG islands, which is significantly higher than for non-ubi-PLSs (45%, 11,603 of 25,705, Fisher’s exact test *p*-value < 2.2 × 10^-16^). Moreover, 510 out of 25,705 non-ubi-PLSs had zero CpG sites (hence CG content = 0 in **Figure 2A**), while none of ubi-PLSs had zero CpG sites. The high CG content of ubi-PLSs suggests that these promoters would support strong and ubiquitous transcription programs.

**Figure 2.**
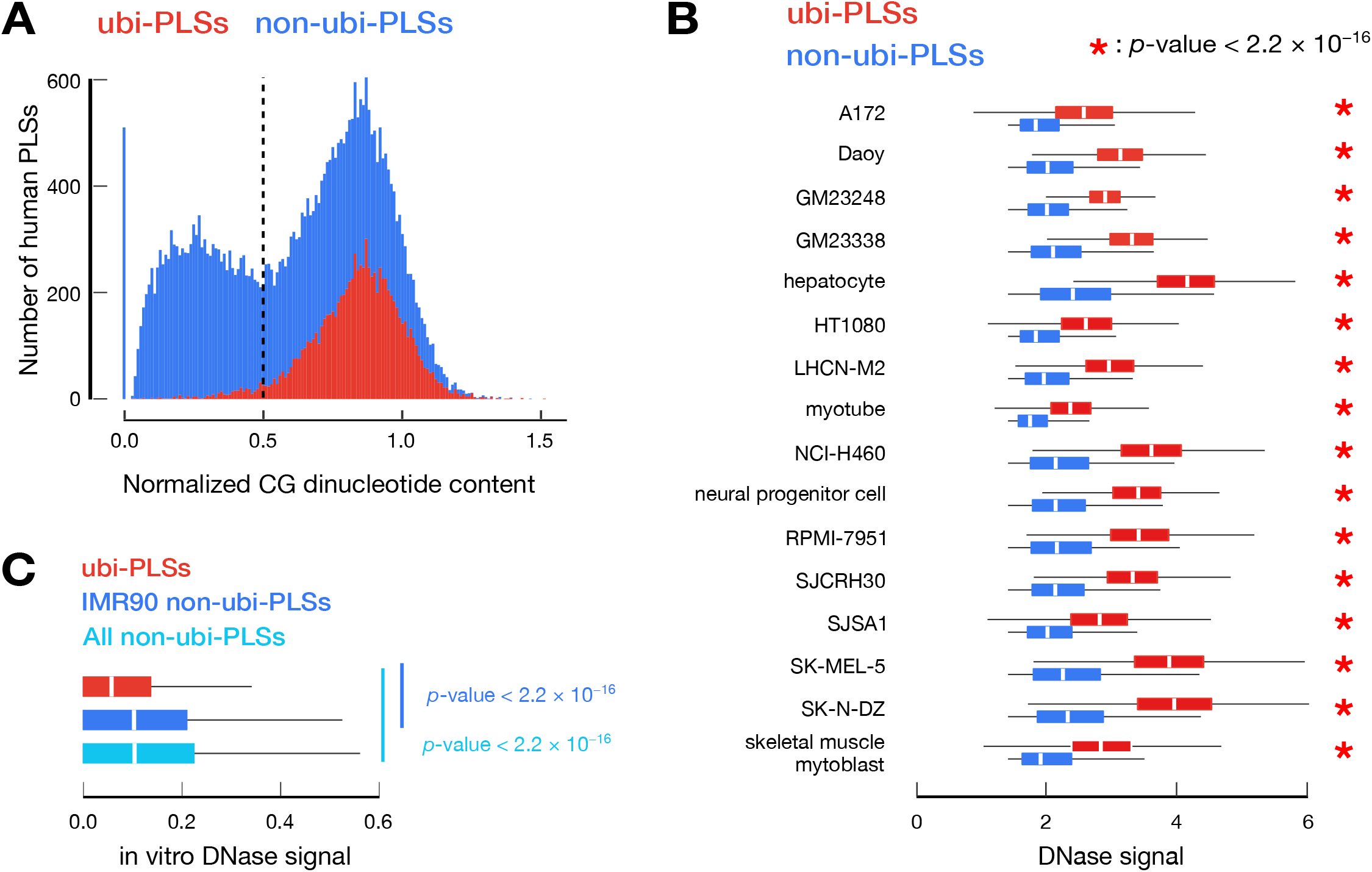
ubi-PLSs are enriched in CG dinucleotides and are open chromatin in cells but closed chromatin in vitro. **A.** ubi-PLSs have higher CG content than non-ubi-PLSs. Histograms show that a vast majority of ubi-PLSs (red) have high CG dinucleotide content (> 0.5), while non-ubi-PLSs (blue) show a bimodal distribution of CG content. **B.** ubi-PLSs (red) have significantly higher DNase signals than non-ubi-PLSs (blue) defined in the same biosample. Data for 16 biosamples are shown, with each biosample represented by a pair of boxplots. All *p*-values were computed with Wilcoxon rank-sum tests. **C.** ubi-PLSs (red) have significantly lower in vitro DNase signals than non-ubi-PLSs (blue) with the DNase-seq experiment performed on extracted DNA. Wilcoxon rank-sum test: *p*-value < 2.2 × 10^-16^ between ubi-PLSs and non-ubi-PLSs defined in IMR90 cells, and *p*-value < 2.2 × 10^-16^ between ubi-PLSs and all non-ubi-PLSs.

The DNase-seq signals at ubi-PLSs were substantially higher than those at non-ubi-PLSs (**Figure 2B**; 31-67% higher by median; Wilcoxon rank-sum test, *p*-values <2.2 × 10^-16^). We surveyed 16 human biosamples (see **Methods; Supplementary Table 1**) and only used the subset of non-ubi-PLSs defined in each biosample for comparison with all ubi-PLSs. Accordingly, ubi-PLSs had lower nucleosome occupancy levels and more strongly positioned flanking nucleosomes than non-ubi-PLSs based on MNase-seq data in K562 cells (**Supplementary Figure 2**).

However, the DNA sequences of ubi-PLSs do not facilitate their open chromatin status in living cells. Reanalyzing the in vitro DNase-seq data performed using DNA molecules purified from IMR90 cells [14], we found that ubi-PLSs had significantly lower in vitro DNase-seq signals than non-ubi-PLSs (**Figure 2C**). This result is consistent with the high CG content of ubi-PLSs (**Figure 2A**), as high CG is known to facilitate chromatin formation [25]. Thus, the open chromatin status of ubi-PLSs are due to trans-factors in the cell that compete with histones.

Next, we compared the signals of eight histone modifications around ubi-PLSs and non-ubi-PLSs in K562 cells. Consistent with the DNase and MNase results described above, the levels of all histone marks examined were lower at the centers of ubi-PLSs than non-ubi-PLSs (**Supplementary Figure 2**), indicating that ubi-PLSs adopt more open chromatin structures than non-ubi-PLSs. For the flanking regions, especially the downstream regions with respect to the transcriptional direction (indicated by arrows pointing to the right in **Supplementary Figure 2**), the signals of promoter-enriched histone marks (H3K4me3, H3K4me2, H3K9ac, and H3K27ac) and transcription-induced histone mark H3K36me3 were substantially higher for ubi-PLSs than non-ubi-PLSs. However, this is the opposite for the enhancer-enriched histone mark H3K4me1 and the repressive mark H3K27me3 (**Supplementary Figure 2**). Thus, ubi-PLSs have a more conducive epigenetic environment than non-ubi-PLSs for transcriptional output.

We further compared the activity of ubi-PLSs and non-ubi-PLSs using ChIA-PET data [26]. Using four sets of ChIA-PET data (RNAPII and CTCF in HeLa and GM12878), we tested whether ubi-PLSs were more enriched in ChIA-PET loop anchors than non-ubi-PLSs. We observed a significantly higher percentage of ubi-PLSs located at loop anchors than non-ubi-PLSs (Fisher’s exact test, *p*-value < 2.2 × 10^-16^ for all four ChIA-PET datasets; **Supplementary Figure 3A**). Reciprocally, we also observed a higher percentage of loop anchors that overlapped ubi-PLSs than non-ubi-PLSs (**Supplementary Figure 3B**). These results suggest that ubi-PLSs are engaged in more three-dimensional chromatin interactions than non-ubi-PLSs.

### ubi-PLSs have higher transcription levels and display more ubiquitous expression profiles than non-ubi-PLSs

To directly quantify the transcriptional activity of ubi-PLSs, we analyzed RNA-seq and RAMPAGE data in the same 16 biosamples that we analyzed DNase-seq data above (see **Methods**). RNA-seq data revealed that genes with at least one TSS overlapping ubi-PLSs (n = 9,214) had significantly higher expression levels than genes with TSSs overlapping only non-ubi-PLSs in a specific biosample (**Figure 3A**; 3.8-12.4 fold higher by median; Wilcoxon rank-sum test, *p*-values < 2.2 × 10^-16^). Among the 9,214 genes, 5,748-7,817 (62%-85%) were expressed with at least 1 transcript per million reads (TPM) in individual biosamples. When examined at the individual TSS level using RAMPAGE data, TSSs overlapping ubi-PLSs had substantially higher activity levels than TSSs overlapping non-ubi-PLSs in the same biosample (**Figure 3B**; 14.5-fold or higher by median; Wilcoxon rank-sum test, *p*-values < 2.2 × 10^-16^). These results suggest that ubi-PLSs correspond to the promoters of highly expressed genes.

**Figure 3.**
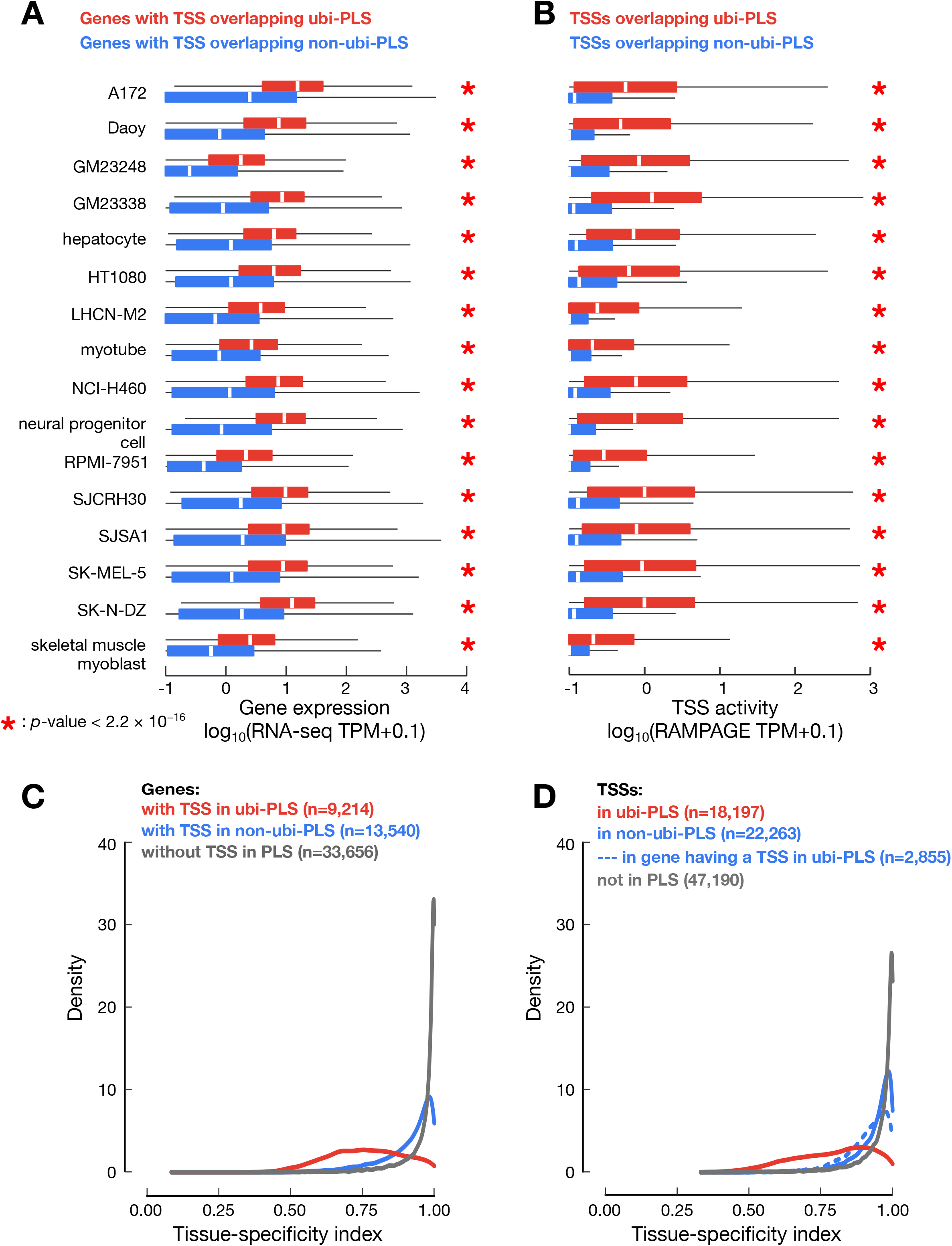
ubi-PLSs are promoters of ubiquitously expressed genes. **A.** Genes whose TSSs overlapped ubi-PLSs (red) are significantly more highly expressed than genes whose TSSs overlapped non-ubi-PLSs (blue) in the same biosample. Expression levels were obtained from RNA-seq data, quantified by TPM (with a pseudocount of 0.1 added to each gene), and plotted in log scale. All *p*-values were computed with Wilcoxon rank-sum tests. **B**. TSSs overlapping ubi-PLSs (red) are significantly more active than TSSs overlapping non-ubi-PLSs (blue) in the same biosample. TSS activities were computed using RAMPAGE data, quantified by TPM (with a pseudocount of 0.1 added to each TSS), and plotted in log scale. All *p*-values were computed with Wilcoxon rank-sum tests. **C.** Distributions of the tissue-specificity index for genes whose TSSs overlapped ubi-PLSs (red), overlap non-ubi-PLS (blue), and do not overlap a PLS (gray). The tissue-specificity index was computed using RNA-seq data across 103 human biosamples (see **Methods**). The three distributions are significantly different (Wilcoxon rank-sum test *p*-value < 2.2 × 10^-16^ for ubi-PLS vs. non-ubi-PLS or vs. non-PLS). **D**. Distributions of the tissue-specificity index for TSSs overlapping ubi-PLSs (red), TSSs of the genes without a ubi-PLS overlapping TSS but with at least one TSS overlapping a non-ubi-PLS (solid blue), other TSSs of the genes with at least one TSS overlapping a ubi-PLS (dashed blue), and TSSs not overlapping a PLS (gray). All the distributions are significantly different (Wilcoxon rank-sum test, *p*-values < 2.2 × 10^-16^ for TSSs overlapping ubi-PLS vs. each of the other three groups of TSSs).

Because promoters with high CG dinucleotide content tend to be ubiquitously expressed, and because ubi-PLSs are enriched in CG dinucleotides, we next evaluated the tissue specificity of ubi-PLSs by surveying 103 ENCODE biosamples with both RNA-seq and RAMPAGE data (for evaluation at the gene and TSS levels, respectively). Using a tissue-specificity index [27], genes with one or more TSSs overlapping ubi-PLSs were substantially less tissue-specific than genes with one or more TSSs overlapping non-ubi-PLSs but none overlapping ubi-PLSs (**Figure 3C**; Wilcoxon rank-sum test, *p*-value < 2.2 × 10^-16^; median = 0.76 and 0.94, respectively). Both of these groups were more tissue-specific than the genes whose TSSs did not overlap any cCRE-PLSs (**Figure 3C**; Wilcoxon rank-sum test, *p*-value < 2.2 × 10^-16^; median = 0.99 for genes with no TSSs overlapping PLS), presumably because genes in the latter group were not expressed in the large number of biosamples assayed by ENCODE, consistent with their high tissue specificity. With the tissue-specificity index [27] computed using RAMPAGE data for individual TSSs, the TSSs overlapping ubi-PLSs showed much lower tissue specificity than the TSSs overlapping non-ubi-PLSs (**Figure 3D**; Wilcoxon rank-sum test, *p*-value < 2.2 × 10^-16^; median = 0.81 and 0.96, respectively). A small number of TSSs (n = 2,855) overlapping non-ubi-PLSs but belonged to genes that had other TSSs overlapping ubi-PLSs; these TSSs were significantly less tissuespecific than the remaining 22,263 TSSs overlapping non-ubi-PLSs (**Figure 3D**; Wilcoxon ranksum test, *p*-value < 2.2 × 10^-16^; median = 0.93 and 0.96, respectively). Thus, genes and TSSs associated with ubi-PLSs are broadly expressed across many biosamples.

### ubi-PLSs contain multiple TSSs and are depleted of TATA-box and narrow-peak promoters

While most human promoters have high CpG content but are depleted in TATA boxes, a subset of promoters contain a TATA box approximately 30 nt upstream of the TSS. These two classes of promoters exhibit ubiquitous and tissue-specific expression, respectively, with expressed TSSs showing distinct sequencing-read profiles (called promoter shapes) [28,29]. As shown above, ubi-PLSs mostly correspond to high-CG promoters with ubiquitous expression. We further found that most ubi-PLSs contained multiple GENCODE TSSs while most non-ubi-PLSs contained only one TSS (**Figure 4A**; Wilcoxon rank-sum test, *p*-value < 2.2 × 10^-16^). Thus, we proceeded to analyze their promoter shapes.

**Figure 4.**
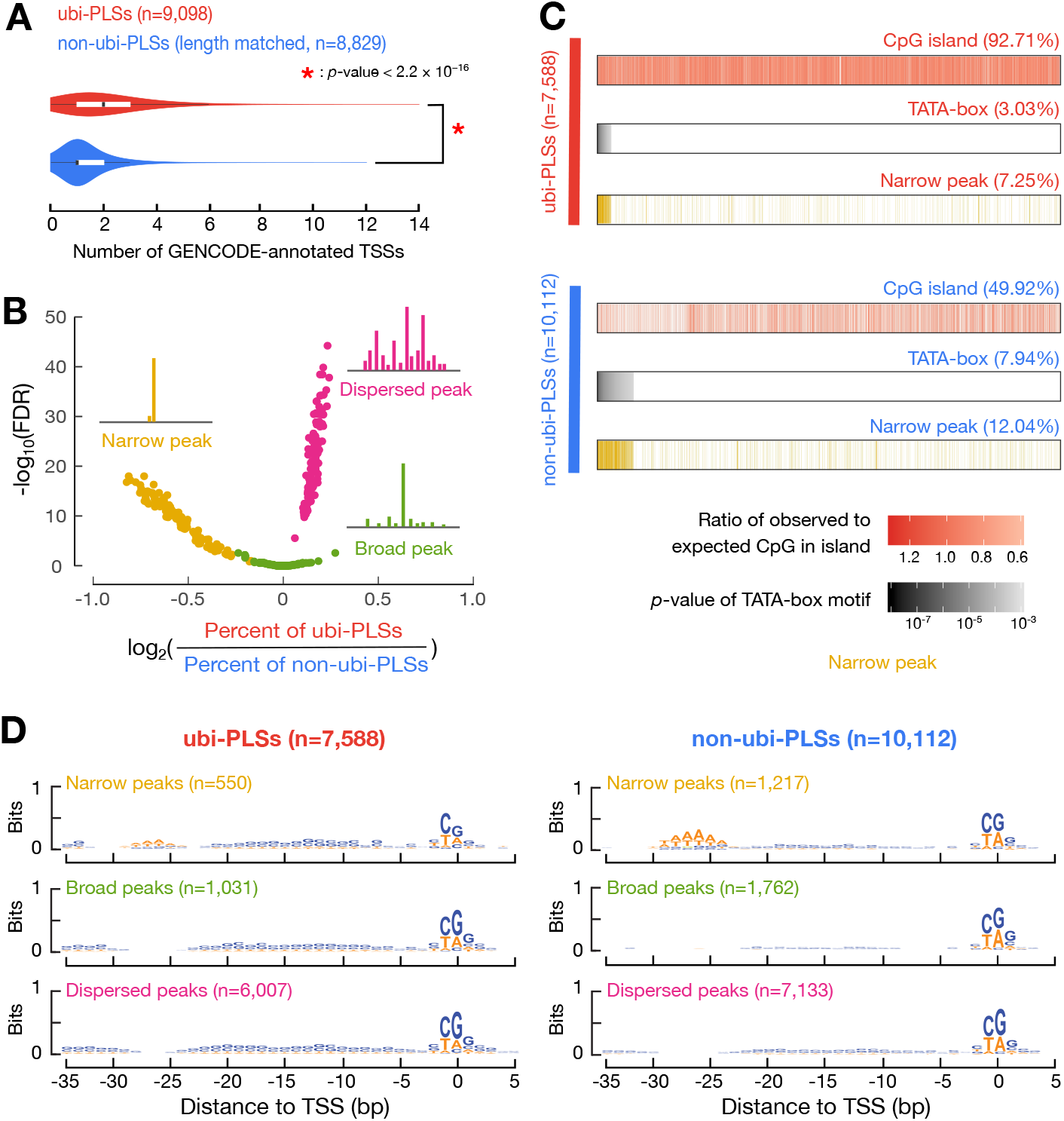
The promoter features of ubi-PLSs. **A.** ubi-PLSs (red) tend to have multiple GENCODE-annotated TSSs while non-ubi-PLSs (blue) normally contain one TSS. A subset of non-ubi-PLSs were used to match the length distribution of ubi-PLSs. The *p*-value was computed with a Wilcoxon rank-sum test. **B.** ubi-PLSs are enriched in dispersed peaks, while non-ubi-PLSs are enriched in narrow peaks. The three types of promoter peak shape (narrow, broad, dispersed) were defined using RAMPAGE data in 115 biosamples (see **Methods**). Each RAMPAGE dataset is represented by three points, one yellow, one red, and one green, plotting the enrichment for narrow, dispersed, and broad peaks, respectively. For each point, the x-axis plots the log-ratio of the percentage of ubi-PLSs over the percentage of non-ubi-PLSs assigned to a peak shape in that RAMPAGE dataset, and the y-axis plots the significance of the enrichment in that peak shape (-log of the Fisher’s exact test *p*-values after FDR correction). **C.** ubi-PLSs are more likely to overlap CpG islands and less likely to have TATA-boxes than non-ubi-PLSs. Two groups of bars with columns depict ubi-PLSs and non-ubi-PLSs, respectively. The top bar in each group shows the enrichment of CpG islands, the middle bar shows the enrichment of TATA-boxes, and the bottom bar shows whether a PLS is a narrow peak. The ubi-PLSs and non-ubi-PLSs (individual columns in each bar) are sorted from left to right by the −log10(*p*-value of the TATA-box motif). **D.** Sequence logos of ubi-PLSs and non-ubi-PLSs with narrow, broad, and dispersed peaks, respectively.

Using RAMPAGE data in each of 115 biosamples, we classified promoters into three categories using a previous definition [29] (**Supplementary Figure 4**; see **Methods**): narrow peaks (a narrow RAMPAGE peak with a single summit that contains most of the reads), broad peaks (a broad RAMPAGE peak with a single summit that contains most of the reads), and dispersed peaks (RAMPAGE peaks without a single summit that contains most of the reads). For example, the *SNRPD1* gene had multiple TSSs overlapping a ubi-PLS, and it contained a dispersed peak that was transcribed in both K562 and GM12878 cells (**Supplementary Figure 5A**). In sharp contrast, the *CALB1* gene contained one TSS in a non-ubi-PLS showing a narrow peak shape that was transcribed in K562 cells but not in GM12878 cells (**Supplementary Figure 5B**). In comparison with non-ubi-PLSs, ubi-PLSs were moderately but significantly enriched in dispersed-peak promoters while depleted in narrow-peak promoters in most of the 115 biosamples (**Figure 4B**).

The vast majority (92.71%) of ubi-PLSs overlapped CpG islands, while only half of non-ubi-PLSs overlapped CpG islands (**Figure 4C**). In contrast, the percentage of non-ubi-PLSs that had a TATA-box was twice that of ubi-PLSs (7.94% versus 3.03%; **Figure 4C**). The TATA-box preference was well-matched with promoter peak shape, with TATA promoters corresponding to narrow peaks (**Figure 4C**). When ubi-PLSs and non-ubi-PLSs were divided into three groups according to their promoter shapes, narrow peaks showed a prominent TATA-box at the −30 nt position while broad and dispersed peaks were depleted of any sequence motif at this position; this pattern was observed for both ubi-PLSs and non-ubi-PLSs (**Figure 4D**). Nevertheless, the narrow peaks in non-ubi-PLSs showed a stronger TATA motif than the narrow peaks in ubi-PLSs, while each promoter shape in ubi-PLSs were more enriched in C and G nucleotides than the corresponding promoter shape in non-ubi-PLSs (**Figure 4D**). Our results are consistent with the previously reported enrichment of high-CG promoters in dispersed peaks and TATA promoters in narrow peaks [28,29]. In summary, ubi-PLSs and non-ubi-PLSs have distinct promoter features— ubi-PLSs tend to have multiple TSSs and are enriched in dispersed peaks while depleted of narrow peaks, and most ubi-PLSs overlap CpG islands and are unlikely to have a TATA-box.

### ubi-PLSs are enriched in the motifs of ubiquitously expressed transcription factors and the promoters preferentially responsive to EMSY and MLL3

We scanned the two sets of cCRE-PLSs against the transcription factor motifs in the JASPAR database (see **Methods**) and found that ubi-PLSs had significantly more motif sites than non-ubi-PLSs; collectively, these motif sites covered more base pairs in ubi-PLSs than in non-ubi-PLSs (**Figure 5A**). Two different sets of transcription factors corresponded to these motif sites, with the ETV and ELK families among 38 transcription factors showing the strongest enrichments for ubi-PLSs and the HNF, GATA, and AP1 families among 119 transcription factors showing the strongest enrichments for non-ubi-PLSs (**Figure 5B, Supplementary Table 4**). Some of these enriched transcription factors were further supported by enrichments in binding signals using available ChIP-seq data in four human cell lines (HepG2, H1, K562, and GM12878; **Supplementary Figure 6**). The ChIP-seq signals were higher at ubi-PLSs than at non-ubi-PLSs for all transcription factors, regardless of their preferences for one set of cCRE-PLSs over the other (**Supplementary Figure 7**). The transcription factors with motif enrichment in ubi-PLSs showed ubiquitous expression profiles computed using the aforementioned RNA-seq data in 103 ENCODE biosamples (see **Methods**), while the transcription factors with motif enrichment in non-ubi-PLSs showed highly tissue-specific expression profiles (**Figure 5C**). Moreover, the transcription factors with motif enrichment in ubi-PLSs were expressed at higher levels (measured by RNA-seq in the same 16 samples as in **Figure 3A**) than the transcription factors with motif enrichment in non-ubi-PLSs (**Supplementary Figure 8**). In summary, ubi-PLSs are bound by more transcription factors and are bound more strongly than non-ubi-PLSs, and the transcription factors that regulate ubi-PLSs are more widely expressed across cell types than the transcription factors that regulate non-ubi-PLSs. These findings are highly consistent with the expression profiles of the two sets of cCRE-PLSs as described in previous sections.

**Figure 5.**
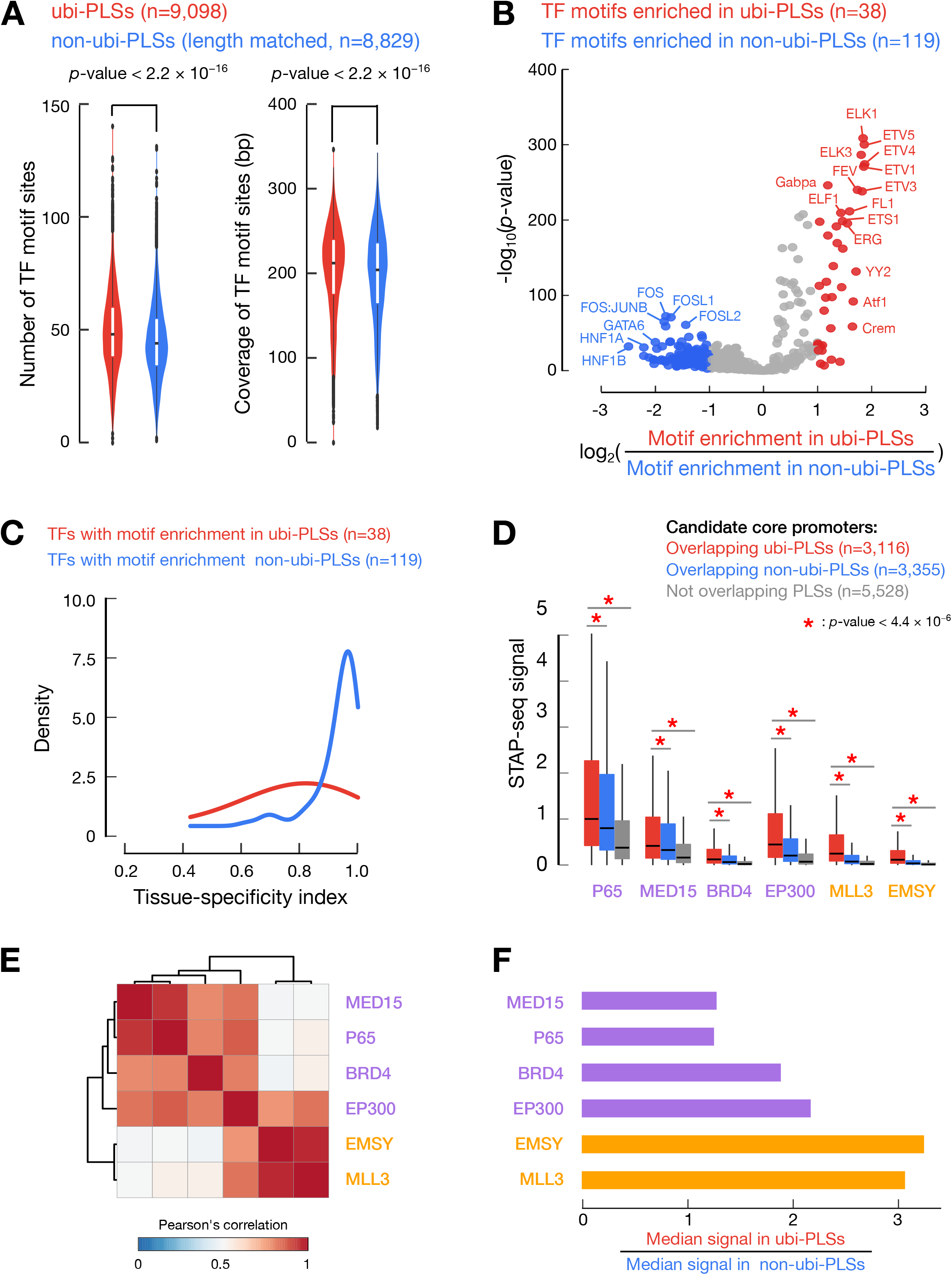
ubi-PLSs have more transcription factor binding sites than non-ubi-PLSs, and the two sets of PLSs are enriched in different transcription factor motifs. **A.** Violin plots compared the number of transcription factor binding sites (left) and the total number of genomic positions covered by transcription factor binding sites (coverage, right) between ubi-PLSs (red) and non-ubi-PLSs (blue). Non-ubi-PLSs were downsampled to match the length distribution of ubi-PLSs for a fair comparison. All *p*-values were computed with Wilcoxon ranksum tests. **B.** The volcano plot shows the enriched transcription factor motifs for ubi-PLSs vs. non-ubi-PLSs. Each dot is a transcription factor motif. The x-axis is the log fold change of the transcription factor motif enrichment in ubi-PLSs over non-ubi-PLSs, and the y-axis depicts Fisher’s exact test *p*-value of the enrichment. Transcription factors whose motifs are enriched in ubi-PLSs are in red, while transcription factors whose motifs are enriched in non-ubi-PLSs are in blue. TF: transcription factor. **C.** Transcription factors that prefer to bind ubi-PLSs are less tissue-specific than the transcription factors that prefer to bind non-ubi-PLSs. The two groups of transcription factors are defined by their enriched motifs in panel **B**, and the tissue-specificity index was calculated using RNA-seq data across 103 samples as in **Figure 3C**. TF: transcription factor. **D.** Boxplots show the activities (STAP-seq signals) of the promoter candidates that overlapped ubi-PLSs (red), overlapped non-ubi-PLSs (blue), or did not overlap PLSs (gray). STAP-seq data on six transcriptional cofactors were available and the colors of the cofactors represent their preference for regulating TATA-box (purple) or CG-rich (orange) promoters. All *p*-values were computed with Wilcoxon rank-sum tests. **E.** Hierarchical clustering of the six cofactors based on Pearson’s correlation of STAP-seq signals across the 6,471 promoter candidates overlapping PLSs. **F.** Ratios of the median STAP-seq signals for promoter candidates overlapping ubi-PLSs versus those overlapping non-ubi-PLSs.

We further assessed the promoter activities of the two sets of cCRE-PLSs using the STAP-seq data from a high-throughput promoter activity assay [30]. The STAP-seq data were available for six transcriptional cofactors with distinct preferences for promoter types—MED15, P65, BRD4, and EP300, which prefer TATA-box promoters, and EMSY and MLL3, which prefer CpG-island promoters [30]. For all six cofactors, the candidate core promoters that overlapped ubi-PLSs had significantly higher STAP-seq signals than those that overlapped non-ubi-PLSs, which were further higher than those that did not overlap either group of cCRE-PLSs (**Figure 5D;** Wilcoxon rank-sum test, *p*-values < 4.4 × 10^-6^). This result is consistent with our earlier results on the higher expression levels for genes and TSSs that overlapped ubi-PLSs across all biosamples analyzed (**Figure 3**). The promoter activity profiles for the six cofactors clustered into two groups when only the candidate core promoters were used (**Figure 5E**), in the same way as reported previously with all candidate core promoters [30]. The two cofactors that were known to prefer CpG-island promoters over TATA-box promoters—EMSY and MLL3—exhibited the highest ratios of median signals among the candidate core promoters that overlapped ubi-PLSs over the median signals among the candidate core promoters that overlapped non-ubi-PLSs (**Figure 5F**). This result is also consistent with our earlier results on the enrichment of ubi-PLSs in CpG-island promoters (**Figure 4**). Thus, ubi-PLSs have higher promoter activities than non-ubi-PLSs, and they prefer different transcriptional cofactors.

### ubi-PLSs are highly conserved between human and mouse

To explore the evolutionary conservation of ubi-PLSs between human and mouse, we defined mouse ubi-PLSs in the same way as human ubi-PLSs (see **Methods**). Mouse rDHSs with high DNase signal in at least 90 biosamples (out of a total of 94 biosamples with DNase-seq data) were defined as ubi-rDHSs (**Figure 6A**; n = 13,247). Similar to the human ubi-rDHSs, the majority of mouse ubi-rDHSs were TSS-proximal (60% were PLS and 22% were pELS; **Figure 6B**); in total, there were 7,907 mouse ubi-PLSs (**Supplementary Table 2D**). These ubi-PLSs overlapped 12,759 GENCODE M18 TSSs, which belonged to 8,058 genes (**Supplementary Table 2E, F**).

**Figure 6.**
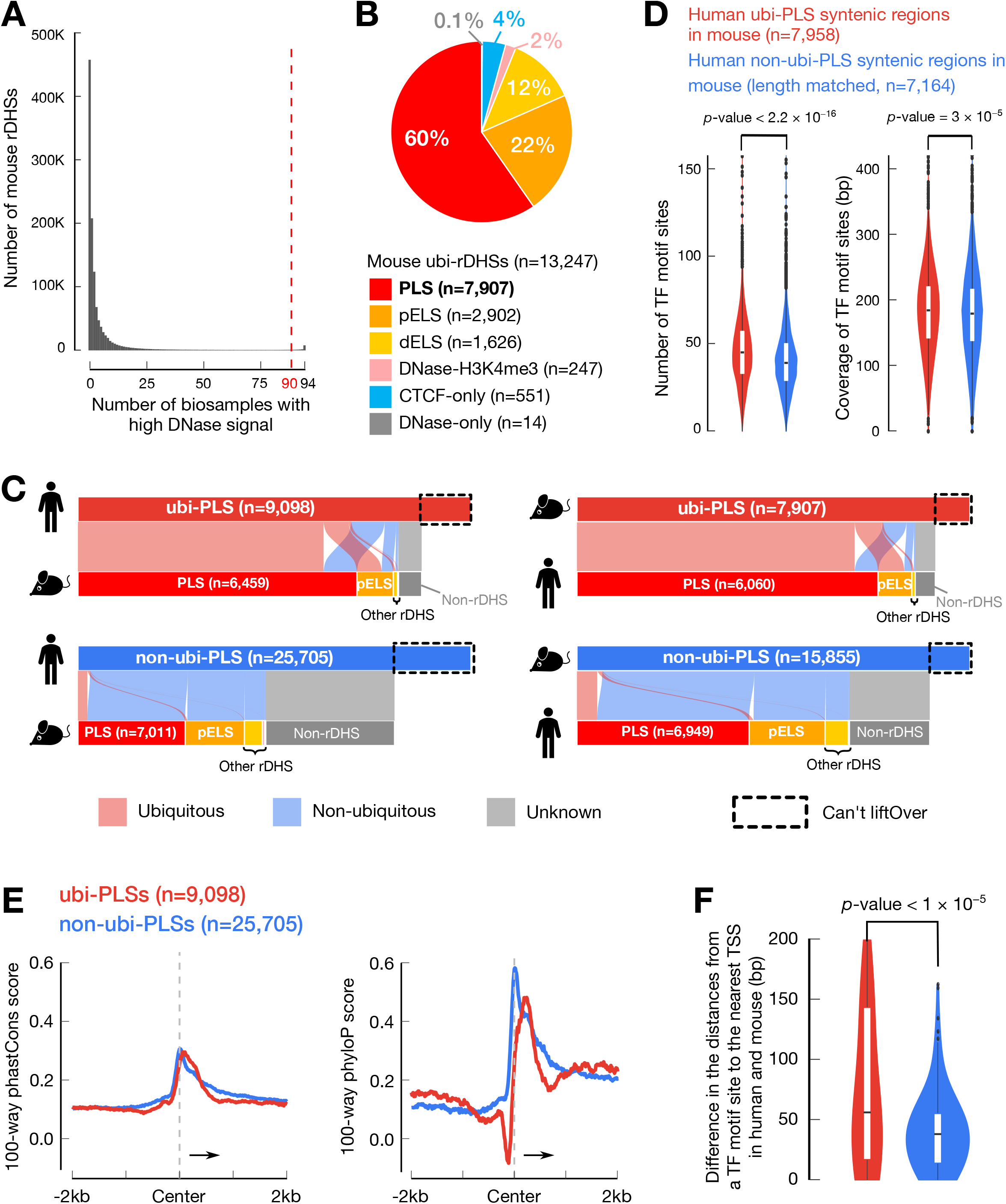
Human ubi-PLSs are conserved in mouse. **A.** Definition of mouse ubi-rDHSs. A histogram shows the number of mouse rDHSs that have high DNase signals in a certain number of biosamples. rDHSs that have high DNase signals in 90 or more biosamples are defined as ubi-rDHSs in mouse. **B**. The pie chart shows that 82% of mouse ubi-rDHSs are PLSs or pELSs. The category of ubi-rDHSs are as in **Figure 1B**. **C.** Most human ubi-PLSs are also mouse ubi-PLSs and vice versa. The two alluvial plots on the left show the syntenic regions of human PLSs in the mouse genome, and the two alluvial plots on the right show the syntenic regions of mouse PLSs in the human genome. The color of a ribbon indicates whether a PLS is ubiquitous (pink) or not (light blue) in the other genome, while a gray ribbon indicates that although some PLSs can be lifted over to the other genome, they are no longer rDHSs in that other genome. **D.** Human ubi-PLSs maintain their higher density and diversity of transcription factor binding sites in the mouse genome. As in **Figure 4A**, mouse regions that are lifted over from human ubi-PLSs (red) have more transcription factor binding sites and higher transcription factor binding site coverage than the mouse regions lifted over from human non-ubi-PLSs (blue). All *p*-values were computed with Wilcoxon rank-sum tests. TF: transcription factor. **E.** Average 100-way phastCons scores (left) and phyloP scores (right) of the ± 2 kb genomic regions centered on human ubi-PLSs (red) and non-ubi-PLSs (blue). **F.** Violin-box plots show the distributions of the difference between human and mouse in the distance from transcription factor motif sites in ubi-PLSs (red) and non-ubi-PLSs (blue) to the nearest TSS. The Wilcoxon rank-sum test *p*-value is shown. TF: transcription factor.

The vast majority of ubi-PLSs and non-ubi-PLSs in both species could be lifted over to the reciprocal genome (see **Methods**), with a slightly higher mapping rate for ubi-PLSs (87% from human to mouse and 91% from mouse to human) than for non-ubi-PLSs (81% from human to mouse and 90% from mouse to human; **Figure 6C**). Most of the syntenic regions of ubi-PLSs in the other genome were still ubi-PLSs (63% of the human ubi-PLSs and 71% of the mouse ubi-PLSs). In sharp contrast, much lower percentages of the syntenic regions of non-ubi-PLSs remained non-ubi-PLSs in the other species (25% of human non-ubi-PLSs and 39% of mouse non-ubi-PLSs); they became pELSs or were no longer rDHSs (**Figure 6C**). Thus, ubi-PLSs are much more evolutionarily conserved than non-ubi-PLSs in terms of synteny and ubiquity of chromatin accessibility.

Like human ubi-PLSs (**Figures 5A**), the syntenic regions of human ubi-PLS in mouse had significantly more motif sites than the syntenic regions of human non-ubi-PLS in mouse, and collectively, these motif sites covered more base pairs in the syntenic regions of ubi-PLSs than non-ubi-PLSs (**Figure 6D**).

We further examined the sequence conservation of the ± 2 kb region centered on the two sets of cCRE-PLSs using two metrics for evolutionary conservation, phastCons and phyloP [31–33]. ubi-PLSs are less conserved than non-ubi-PLSs in the small window immediately upstream of the cCRE-PLS center as revealed by phyloP, which has a higher resolution than phastCons (**Figure 6E**). This small window corresponds to the core promoter where most transcription factors bind. We described above that ubi-PLSs had more transcription factor motif sites than non-ubi-PLSs (**Figures 5A**, **6D**), and we asked whether these motif sites maintained their positions between the two species. We found that the motif sites in ubi-PLSs tended to differ more greatly in their distances to the nearest TSS between human and mouse than the motif sites in non-ubi-PLSs (**Figure 6F**). Thus, the turn over of transcription factor binding sites may be a cause for the lower sequence conservation in the core promoter regions that correspond to ubi-PLSs than those of non-ubi-PLSs.

## Discussion

We performed systematic analysis on 9,098 human promoters and 7,907 mouse promoters that had open chromatin in more than 95% of the human and mouse biosamples tested. Compared with non-ubi-PLSs, ubi-PLSs have seven striking genomic and epigenomic features in addition to having ubiquitously open chromatin. First, they have high CG content and lack a TATA box (**Figures 2A**, **4C**). Second, they mostly belong to housekeeping genes, being highly enriched in GO categories of metabolism and biosynthesis and highly depleted in GO categories of signal transduction and stimulus detection (**Figure 1E**, **F**). Accordingly, they are also highly enriched in essential genes (**Figure 1G, H**). Third, these ubi-PLSs show high levels of open chromatin (**Figure 2B**), active histone modifications, well-positioned flanking nucleosomes (**Supplementary Figure 2**), and high levels of chromatin interactions (**Supplementary Figure 3**). Fourth, they are highly expressed (at both the gene and TSS levels) across cell and tissue types (**Figure 3**). Fifth, transcription tends to fire from multiple positions in ubi-PLSs, hence ubi-PLSs are enriched in the dispersed-peak promoter shape and depleted in the sharp-peak promoter shape (**Figure 4**). Sixth, they are likely regulated by a distinct set of transcription factors (based on motif and ChIP-seq peak enrichments), which also tend to be ubiquitously expressed (**Figure 5**, **Supplementary Figure 6**), and transcriptional cofactors EMSY and MLL3, which prefer CpG-island promoters (**Figure 5**). Seventh, they are highly conserved between human and mouse at the synteny level, and most of them are ubi-PLSs in both species; however, they are not as conserved at the sequence level, with a high turnover of transcription factor motif sites (**Figure 6**).

Collectively, these genomic and epigenomic features point to a highly consistent model (**Figure 7**) for the transcriptional regulation of roughly nine thousand genes that are expressed in most cell types. With high CG content, the DNA of ubi-PLSs would be bound by nucleosomes in vitro [14] (**Figure 2C**), indicating that it is not the DNA sequences of the ubi-PLSs that have the inherent ability to keep them open chromatin across most cell types. Rather, it is more likely that the occupancy by transcription factors, especially the transcription factors that are expressed in most cell types, competes with histone proteins for accessing the DNA and keeps the ubi-PLSs in the open chromatin state. Indeed, the high ChIP-seq signals of most transcription factors at ubi-PLSs (**Supplementary Figure 7**) support their high occupancy on the ubi-PLSs.

**Figure 7.**
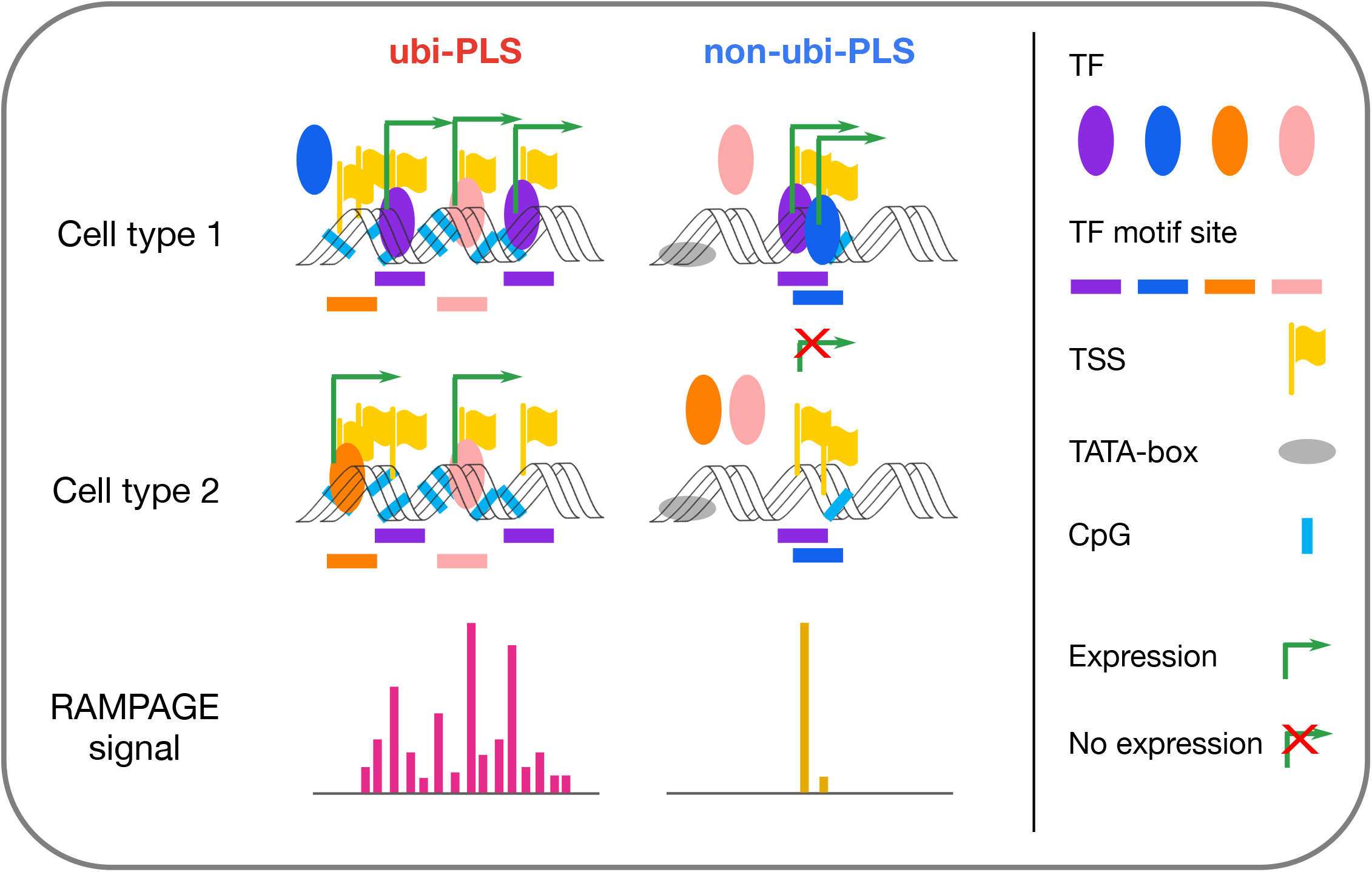
A schematic model of transcriptional activation at ubi-PLSs and non-ubi-PLSs. Ubi-PLSs tend to have high CG content, lack a TATA-box, be bound by multiple transcription factors, and initiate transcription from dispersed genomic positions. They recruit different transcription factors to initiate transcription in different cell types, maintaining high and ubiquitous expression. In contrast, non-ubi-PLSs are more likely to have lower CG content, contain a TATA-box, be bound by fewer transcription factors, and initiate transcription from a specific genomic position. They are expressed in a few cell types and controlled by a select group of transcription factors. TF: transcription factor; TSS: transcription start site.

Typically, multiple transcription factors bind to a promoter and regulate its transcription, and two broad classes of models have been proposed for the mechanisms of gene regulation: the regulatory grammar model [34,35] and the flexible billboard model [36,37]. The regulatory grammar model states that a specific syntax of transcription factor binding (e.g., the relative orientation and distance between neighboring sites) is required for the co-regulation to occur, while the flexible billboard model states that the composition of the bound transcription factors, but not so much their relative orientations and positions, determines the co-regulation. We found that between human and mouse, most syntenic regions of ubi-PLSs in one species were also ubi-PLSs in the other species, although their DNA sequences were less conserved, with movements of transcription factor motif sites (**Figure 6**). Our results suggest that ubi-PLSs are more likely to adopt the flexible billboard model—although the transcription factor sites can change their positions between human and mouse, the promoters remain functional as ubi-PLSs in both species.

Our findings of ubi-PLSs are highly consistent with earlier work based on transcriptome data [28,29]. An earlier study used CAGE data to define different promoter shapes and then analyzed the sequence features for the different promoter shapes, showing that broad-peak promoters were high in CG and lacked a TATA-box [28]. We started with chromatin accessibility data and defined a set of promoters that had open chromatin in most biosamples, and then we showed that ubi-PLSs were enriched in broad-peak, high CG promoters. The earlier study reported that the high CG promoters were more rapidly evolving in mammals than TATA-containing promoters. In agreement with these earlier findings, we found lower sequence conservation in ubi-PLSs than non-ubi-PLSs between human and mouse (**Figure 6**). Thus, our results added chromatin and epigenetic data to the previous knowledge of promoter types defined using transcriptome data, completing our understanding of how promoters are regulated.

High-throughput techniques such as massively parallel reporter assays (MPRA) are increasingly used to dissect the regulatory mechanisms of regulatory elements [38–41]. Some studies insert transcription factor motif sites into synthetic constructs, while other studies eliminate existing motif sites in genomic sequences, and then the impact of these changes is measured by a reporter. One recurrent finding is that the degree of transcription factor occupancy (often represented by the number of motif sites) is one of the best predictors for the reporter expression [38–41]. Our finding of larger numbers of motif sites in ubi-PLSs and their higher expression levels than non-ubi-PLSs agrees with the earlier findings of MPRA studies [39]. Furthermore, we found a large enrichment of ubi-PLSs in bidirectional promoters, which agrees with the earlier finding that bidirectional promoters were more active than unidirectional promoters in driving reporter expression [39]. We found GABPA, E2F2/3, and YY1/2 to be among the 38 transcription factors whose motifs were enriched in ubi-PLSs, and these motifs were found earlier to be enriched in bidirectional promoters, especially GABPA [42]. We found that ubi-PLSs were enriched in the motifs of ubiquitously expressed transcription factors, including the ETV and ELK families, and these motifs were also found to be enriched in the promoters of ubiquitously expressed genes [43].

In summary, we have performed extensive analyses on roughly nine thousand human promoters that are open chromatin in more than 95% of the biosamples. Our analyses showed that these promoters are likely regulated by a common mechanism at the center of which is a small set of widely expressed transcription factors. These promoters are mostly syntenic between human and mouse and are conserved in their ubiquitously expressed promoter function. They are essential for maintaining the high-level transcription of a set of genes required for the normal function of most cell types.

## Methods

### Definition of ubi-rDHSs and ubi-PLSs (Figure 1A-C, Figure 6A, 6B, and Supplementary Table 2)

To define ubi-rDHSs, we started with rDHSs defined by the ENCODE consortium [12] (**Supplementary Table 1A, B**) and calculated the DNase signals (expressed as a Z-score) for each rDHS in all biosamples (517 in total for human and 94 in total for mouse). Following the method by the ENCODE consortium [12], a high signal is defined as Z-score > 1.64, with the threshold of 1.64 corresponding to a *p*-value of 0.05 in a one-sided Z-test. We defined ubi-rDHSs as having high DNase signals in 500 or more human biosamples or 90 or more mouse biosamples. We arrived at 15,989 human ubi-rDHSs (from 2,157,387 human rDHSs) and 13,247 mouse ubi-rDHSs (from 1,192,301 mouse rDHSs). The remaining rDHSs were called non-ubi-rDHSs (**Figure 1A, Figure 6A, Supplementary Table 2**).

The ENCODE consortium defined rDHSs with high ChIP-seq signals of the H3K4me3 or H3K37me3 histone modifications or the CTCF transcription factor as cCREs [12]. The cCREs were further classified into groups: having promoter-like signatures (PLSs), having enhancer-like signatures (ELSs; further classified as proximal or distal to an annotated TSS), with high H3K4me3 signals but are distal to annotated TSSs (DNase-H3K4me3), or bound by CTCF but do not have high H3K4me3 or H3K27ac signals (CTCF-only) [12]. We examined ubi-rDHSs by these cCRE categories (**Figure 1B, Figure 6B**), as well as their distance distribution to the nearest TSS (**Figure 1C**).

### Enrichment analysis of genes associated with ubi-PLSs (Figure 1D, G, H, and Supplementary Figure 1)

We examined the types of genes using the GENCODE v24 annotation. We discarded the GENCODE TSSs of transcripts with inactive or uncertain biotypes such as pseudogenes and TEC (to be experimentally confirmed). In total, there are 19,815 and 36,550 GENCODE genes with the protein-coding and other biotypes, respectively. We counted the percentages of genes in each biotype whose TSSs were located only in ubi-PLSs, only in non-ubi-PLSs, in both ubi-PLSs and non-ubi-PLSs (i.e., some TSSs of the gene whose TSSs overlapped ubi-PLSs while some other TSSs of the same gene whose TSSs overlapped non-ubi-PLSs) or not in ubi-PLSs (**Figure 1D**, colored accordingly).

Bidirectional genes (i.e., genes whose TSSs were within 1000 nt of each other and on opposite genomic strands) were defined based on a previous study [44]. We identified 4,755 pairs formed by 8,930 bidirectional genes (some genes belonged to multiple pairs). We counted and performed Fisher’s exact test between bidirectional genes with or without a TSS in ubi-PLSs (**Figure 1D**).

There were 1,874 genes with TSSs overlapping both ubi-PLSs and non-ubi-PLSs (i.e., they have at least one TSS in ubi-PLSs and at least one TSS in non-ubi-PLSs). To investigate whether this number is significantly different from expected, we randomly assigned TSSs to be in ubi-PLSs while maintaining the numbers of TSSs in each gene and then counted the number of genes with TSSs overlapping both ubi-PLSs and non-ubi-PLSs.

The cell-essential gene data were obtained from previous publications [17–19]. We counted the number of cell-essential genes (using the same cut-off as in each of the previous studies) that had ubi-PLSs or non-ubi-PLSs (**Figure 1G**). We also ranked all genes by their essentiality scores [17] and counted the percentage of genes with their TSSs overlapping ubi-PLSs by going down the ranks and examining each nonoverlapping 500-gene window. The percentage of all tested genes with TSSs overlapping ubi-PLSs is plotted as a control horizontal line (**Figure 1H, Supplementary Figure 1**).

### Gene Ontology enrichment analysis (Figure 1E, F, and Supplementary Table 3)

We performed Gene Ontology enrichment analysis using the Panther tool [45] on the 9,214 genes whose TSSs were located in ubi-PLSs (**Supplementary Table 3**). We used Fisher’s exact test with a false discovery rate (FDR) correction and all *Homo sapien* genes as the background (**Supplementary Table 3**). We used the most enriched and depleted (FDR < 1.0 × 10^-10^) GO Biological Process terms to generate two word clouds with the R package *wordcloud* (**Figure 1E, F**).

### Normalized CG content (Figure 2A)

To evaluate whether PLSs are enriched in CG dinucleotides, we used normalized CG content as previously described [24], defined as the ratio of the observed number over the expected number of CG dinucleotides, where the expected number is calculated as [(Fraction of C+Fraction of G)/2]^2^, i.e.,

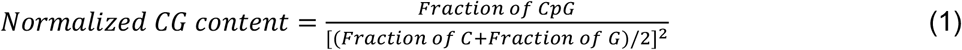

As reported previously [24], promoters show a bimodal distribution for their normalized CG content, with a valley at 0.5 (**Figure 2A**).

### Comparison of DNase, RNA, RAMPAGE, and histone mark levels between ubi-PLSs and non-ubi-PLSs (Figures 2B, 2C, 3A, 3B, and Supplementary Figure 2, 8)

By definition, ubi-PLSs have accessible chromatin in most biosamples while non-ubi-PLSs have accessible chromatin in only a few samples. To compare ubi-PLSs with non-ubi-PLSs at their chromatin and transcriptional levels, we defined non-ubi-PLSs for each biosample as those PLSs that were not ubi-PLSs and yet had a high DNase signal in that sample.

To assess the chromatin accessibility of cCRE-PLSs, we compared the DNase signals between ubi-PLSs and biosample-specific non-ubi-PLSs. DNase-seq data on 16 biosamples were downloaded from the ENCODE portal (ENCODE accessions in **Supplementary Table 1C**) we selected these 16 biosamples because they also had RNA-seq and RAMPAGE data (see below). We performed Wilcoxon rank-sum tests between ubi-PLSs and non-ubi-PLSs (**Figure 2B**).

To assess whether the high chromatin accessibility of ubi-PLSs was due to their intrinsic DNA sequences, we compared the in vitro DNase signals between ubi-PLSs and non-ubi-PLSs using previously published data on DNA purified from IMR90 cells [14] (SRA accession SRX247626). In vitro DNase-seq reads were mapped to GRCh38 using BWA mem [46] and duplicate reads were removed using Picard [47]. Uniquely mappable reads were used to compute signals on ubi-PLSs versus non-ubi-PLSs defined in IMR90 cells or versus all non-ubi-PLSs. We performed Wilcoxon rank-sum tests between ubi-PLSs and non-ubi-PLSs (**Figure 2C**).

To compare the gene expression level and TSS activity between ubi-PLSs and biosample-specific non-ubi-PLSs, we downloaded RNA-seq and RAMPAGE data (ENCODE accessions in **Supplementary Table 1D, E**) in the aforementioned 16 biosamples with DNase-seq data from the ENCODE portal. We then compared the expression levels of genes (using RNA-seq data) and individual TSSs (using RAMPAGE data) associated with ubi-PLSs and non-ubi-PLSs in each of the 16 biosamples. We performed Wilcoxon rank-sum tests between ubi-PLSs and non-ubi-PLSs (**Figure 3A, B**). We also used the RNA-seq signal to compare the expression level of transcription factors with motif enrichment in ubi-PLSs versus non-ubi-PLSs (**Supplementary Figure 8**).

We made meta plots to compare the histone mark ChIP-seq signals between ubi-PLSs and non-ubi-PLSs in K562 cells. For all ubi-PLSs and K562 non-ubi-PLSs, we calculated the average ChIP-seq signals of eight histone marks and the average MNase-seq signal as a measure of nucleosome occupancy. We downloaded ChIP-seq and MNase-seq data from the ENCODE portal (ENCODE accessions in **Supplementary Table 1F, G**) and used the UCSC’s bigWigAverageOverBed tool to calculate the average signal in the ± 2kb window centered on each group of cCRE-PLSs (**Supplementary Figure 2**).

### Chromatin interaction analysis (Supplementary Figure 3)

We downloaded ChIA-PET data [48] from the Gene Expression Omnibus (GEO) with the accession GSE72816. This dataset included RNAPII and CTCF ChIA-PET clusters in HeLa and GM12878 cell lines. We filtered each set of clusters by retaining the ChIA-PET loops that were supported by at least four reads. We then intersected the ChIA-PET loop anchors with ubi-PLSs and biosample-specific non-ubi-PLSs, requiring at least 1 bp overlap. We calculated the percentage of PLSs that overlapped ChIA-PET loop anchors as well as the percentage of ChIP-PET loop anchors that overlapped cCRE-PLSs. Wilcoxon rank-sum tests were performed between ubi-PLSs and non-ubi-PLSs.

### Tissue-specificity index (Figures 3C, 3D, and 5C)

We used a previously defined tissue-specificity index [27] to evaluate the tissue specificity of gene expression and TSS activities, defined as:

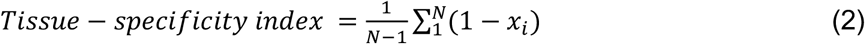

where *N* is the number of biosamples and *x_i_* is the expression profile component normalized by the maximal component value across all biosamples. This tissue-specificity index ranges from 0 to 1, with a higher value indicating a higher degree of tissue specificity.

To calculate the tissue-specificity index, we used RNA-seq and RAMPAGE data from the ENCODE portal (ENCODE accessions in **Supplementary Table 1D, E**), encompassing 103 biosamples for which both types of data were available. Prior to computing the tissue specificity of genes or TSSs, we performed quantile normalization across genes or TSSs in each sample and then applied the above formula on the normalized RNA-seq and RAMPAGE values (**Figure 3C, D**). We also used the RNA-seq data to compute the tissue-specificity index for transcription factors (**Figure 5C**).

### The number of TSSs in cCRE-PLSs (Figure 4A)

We used GENCODE-annotated TSSs (v24, the same version used in ENCODE cCRE definition) [49]. GENCODE-annotated TSSs of transcripts with inactive or uncertain biotypes such as pseudogene and TEC (to be experimentally confirmed) were removed. To remove the impact of the length difference between ubi-PLSs and non-ubi-PLSs, we downsampled non-ubi-PLSs to a subset (n = 8,688) that matched the length distribution of ubi-PLSs. We intersected the genomic coordinates of TSSs with those of PLSs and counted the number of TSSs overlapping each ubi-PLS and non-ubi-PLS in order to draw the violin-box plot (**Figure 4A**). A Wilcoxon rank-sum test was performed between ubi-PLSs and non-ubi-PLSs.

### Definition of RAMPAGE peak shape and calculation of the enrichment of cCRE-PLSs in rPeaks with each shape (Supplementary Figure 4, Figure 4B-D)

We defined the shape of each of the 52,546 representative RAMPAGE peaks (rPeaks) [50] using RAMPAGE data in 115 biosamples downloaded from the ENCODE portal (ENCODE accessions in **Supplementary Table 1E**). Each rPeak was classified as one of three peak shapes according to the flowchart in **Supplementary Figure 4**. Dispersed peaks were defined as rPeaks in which fewer than 50% of the RAMPAGE reads within that peak had their 5’-ends overlapping the region ±2 nt around the peak summit. The remaining rPeaks were further divided into narrow peaks or broad peaks according to peak length (broad peak, > 9 nts; narrow peak, ≤ 9 nts). For each rPeak, we used the same boundary in all biosamples, but the peak summit position and peak shape were determined in each biosample (only for the rPeaks with at least 10 RAMPAGE reads in a sample), which showed small variations across the biosamples.

Using the rPeak shapes defined in 115 biosamples, we computed the enrichment of cCRE-PLSs in rPeaks with each shape type as follows. We calculated the log-ratio between the number of ubi-PLSs and the number of biosample-specific non-ubi-PLSs that overlapped the rPeaks with each shape in each biosample. We further performed a Fisher’s exact test for each biosample and each peak shape, followed by FDR correction of the resulting *p*-value (**Figure 4B**).

For cCRE-PLSs with RAMPAGE peaks in at least one of the 115 biosamples, we tested whether a PLS overlapped a CpG island [51], with CpG island annotations downloaded from the UCSC Genome Browser (hg38, cpgIslandExtUnmasked.txt). We then computed the ratio of the observed over the expected numbers of cCRE-PLSs that overlapped CpG islands (**Figure 4C**). We also tested whether a cCRE-PLS contained a TATA-box by looking for a site for the TATA-box motif within 25-35 bp upstream of the TSS, with the TSS defined by the summit of RAMPAGE peaks. TATA-box sites were predicted using the FIMO algorithm [52] with the parameters --norc --thresh 1e-3 and the TBP position-weight matrix (MA0108.2) from the JASPAR database [53] (**Figure 4C**).

We grouped ubi-PLSs and non-ubi-PLSs according to the shape of rPeaks that they overlapped in the most biosamples, and created sequence logos for each cCRE-PLS sub-group using WebLogo 3 [54] (**Figure 4D**).

### Transcription factor motif and ChIP-seq peak analysis (Figure 5A, 5B, Figure 6D, Supplementary Table 4, and Supplementary Figures 6, 7)

We found that ubi-PLSs were slightly longer than non-ubi-PLSs (median lengths are 349 bp and 295 bp, respectively). To avoid the impact of the length difference on transcription factor motif analysis, we used the downsampled subset of non-ubi-PLSs (n = 8,688) that matched the length distribution of ubi-PLSs as described above for **Figure 5A**. We scanned the two sets of cCRE-PLSs for motif matches using FIMO [52] with the parameters --thresh 1 e-4 and the position-weight matrices downloaded from the JASPAR database [53]. We counted the number of transcription factor motif sites as well as the total number of genomic positions covered by these motif sites in each cCRE-PLS. Wilcoxon rank-sum tests were performed to compare ubi-PLSs and non-ubi-PLSs (**Figure 5A**). We performed the same analysis on the syntenic regions of the human ubi-PLSs and non-ubi-PLSs in the mouse genome (**Figure 6D**). For each transcription factor, we compared the enrichment of its motif between ubi-PLSs and non-ubi-PLSs by both fold enrichment and Fisher’s exact test (**Figure 5B**). Transcription factors with at least two-fold enrichment for their motif sites in ubi-PLS vs. non-ubi-PLS or vice versa were identified (**Supplementary Table 4**).

Furthermore, we extended the transcription factor motif enrichment analysis in a specific biosample by including the transcription factor ChIP-seq data in the same biosample, namely, HepG2, H1, K562, or GM12878 cells. For transcription factors with matching position-weight matrices and ChIP-seq data in one of these four cell lines, we downloaded transcription factor ChIP-seq peaks from the ENCODE portal (ENCODE accessions in **Supplementary Table 1H**). A cCRE-PLS overlapping a transcription factor ChIP-seq peak (by at least half of the peak width) in a cell line and containing a site for the transcription factor motif was considered to be bound by the transcription factor in that cell line. We measured the enrichment by both fold enrichment and Fisher’s exact test in each cell line (**Supplementary Figure 6**).

We made aggregation plots to evaluate the transcription factor ChIP-seq signal on cCRE-PLSs. We downloaded the ChIP-seq signals (bigWig files) of six transcription factors from the ENCODE portal (ENCODE accessions in **Supplementary Table 1H**), with three showing a preference for ubi-PLSs and the other three showing a preference for non-ubi-PLSs (**Figure 5B**). We used UCSC’s bigWigAverageOverBed tool to calculate the average signal across the ± 2kb window centered on cCRE-PLSs (**Supplementary Figure 7**).

### STAP-seq analysis (Figure 5D-F)

We analyzed the promoter activities of ubi-PLSs and non-ubi-PLSs using the STAP-seq data produced using a high-throughput promoter assay [30]. We lifted the genomic coordinates of cCRE-PLSs in GRCh38 to hg19 using the chain files and the liftOver method from the UCSC Genome Browser. We overlapped the 11,979 candidate core promoters tested using the STAP-seq assay in the human HCT116 cell line with our two sets of cCRE-PLSs. We then compared the STAP-seq signals among three groups of candidate core promoters: those that overlapped ubi-PLSs (n = 3,116), those that overlapped non-ubi-PLSs (n = 3,335), and those that did not overlap any cCRE-PLSs (n = 5,528; **Figure 5D**). We performed hierarchical clustering of the cofactors based on Pearson’s correlation of STAP-seq signals across the 6,451 cCRE-PLS-overlapping candidate core promoters (**Figure 5E**). We then calculated the ratio between the median STAP-seq signal of the candidate core promoters that overlapped ubi-PLSs vs. non-ubi-PLSs for each cofactor (**Figure 5F**).

### Evolutionary conservation analysis (Figure 6C, E, F)

We mapped human cCRE-PLSs to the mouse genome (mm10) using the chain files and the liftOver method from the UCSC Genome Browser, with the parameter -minMatch=0.5. We then asked whether these syntenic regions were also cCREs in the mouse genome. A syntenic region in the mouse was considered a cCRE when at least half of the region overlapped a mouse cCRE. These syntenic regions overlapping mouse cCREs were regarded as ubiquitous or not according to the mouse cCREs (**Figure 6C**). We performed a similar analysis to map the mouse cCRE-PLSs to the human genome.

We downloaded the human 100-way phastCons and phyloP signals (bigWig files) from the UCSC Genome Browser. We used UCSC’s bigWigAverageOverBed tool to calculate the average signal across the ± 2 kb windows centered on cCRE-PLSs (**Figure 6E**).

To test whether there is a shift in transcription factor motif sites between human and mouse, we measured the distance from each motif site to the nearest GENCODE-annotated TSS (v24 for human and vM18 for mouse). We only included in this analysis the human cCRE-PLSs that could be lifted over to the mouse genome and their syntenic regions. Wilcoxon rank-sum tests were used to compare the distances between human and mouse (**Figure 6F**).

## Data Availability

All processed datasets are available. cCREs and rDHSs are available at the ENCODE portal (www.encodeproject.org) with the accessions listed in **Supplementary Table 1**. ubi-rDHSs are listed in **Supplementary Table 2**. ChIA-PET datasets were downloaded from GEO under the accession GSE72816. ENCODE RNA-seq, RAMPAGE, ChIP-seq, and MNase-seq experiments are available at the ENCODE portal with accessions listed in **Supplementary Table 1**.

## Supplementary Figure Legends

**Supplementary Figure 1.**
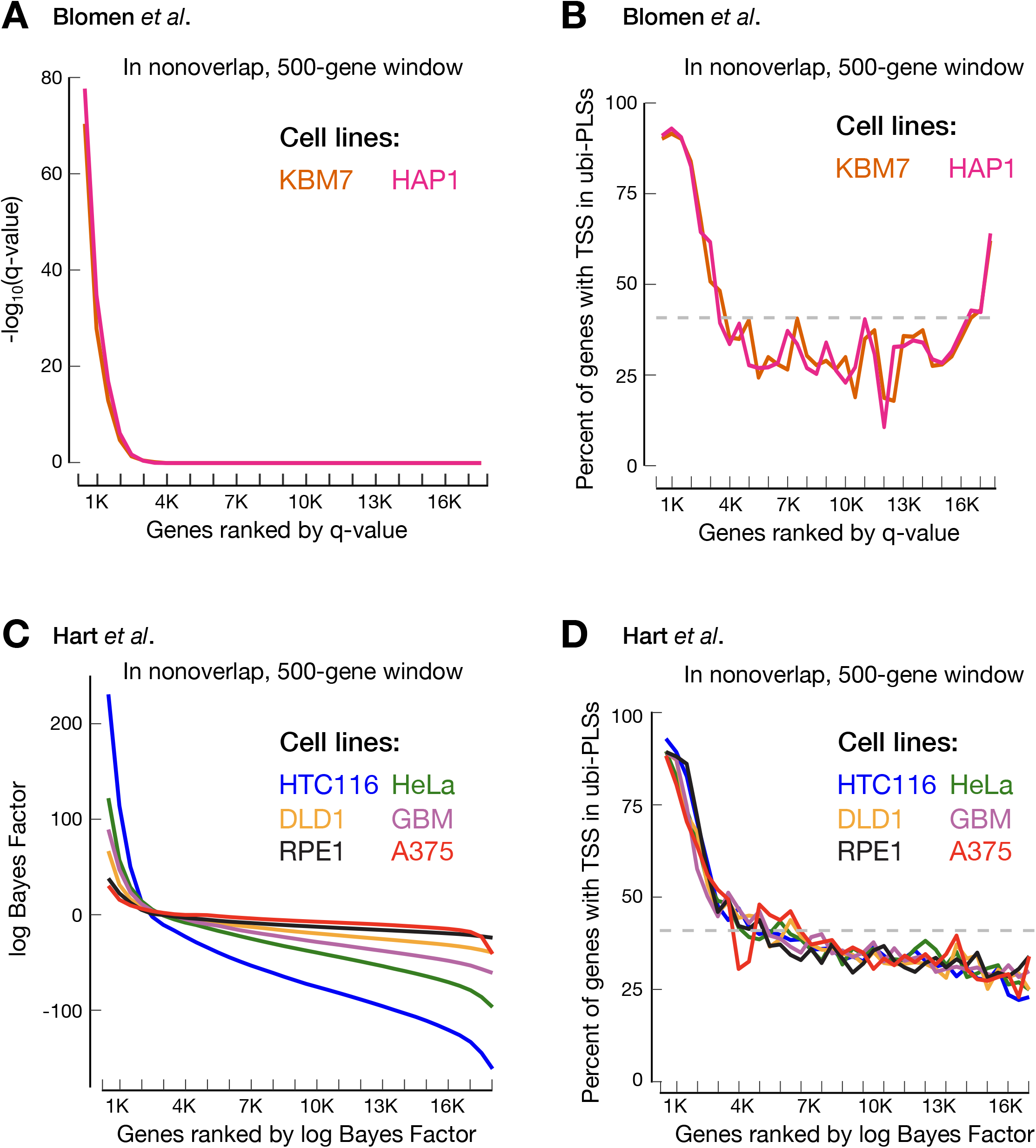
ubi-PLSs are the TSSs of cell-essential genes. Similar to Figure 1H, but with cell-essential genes from the other two studies (Blomen et al. in **A** and **B**, Hart et al. in **C** and **D**).

**Supplementary Figure 2.**
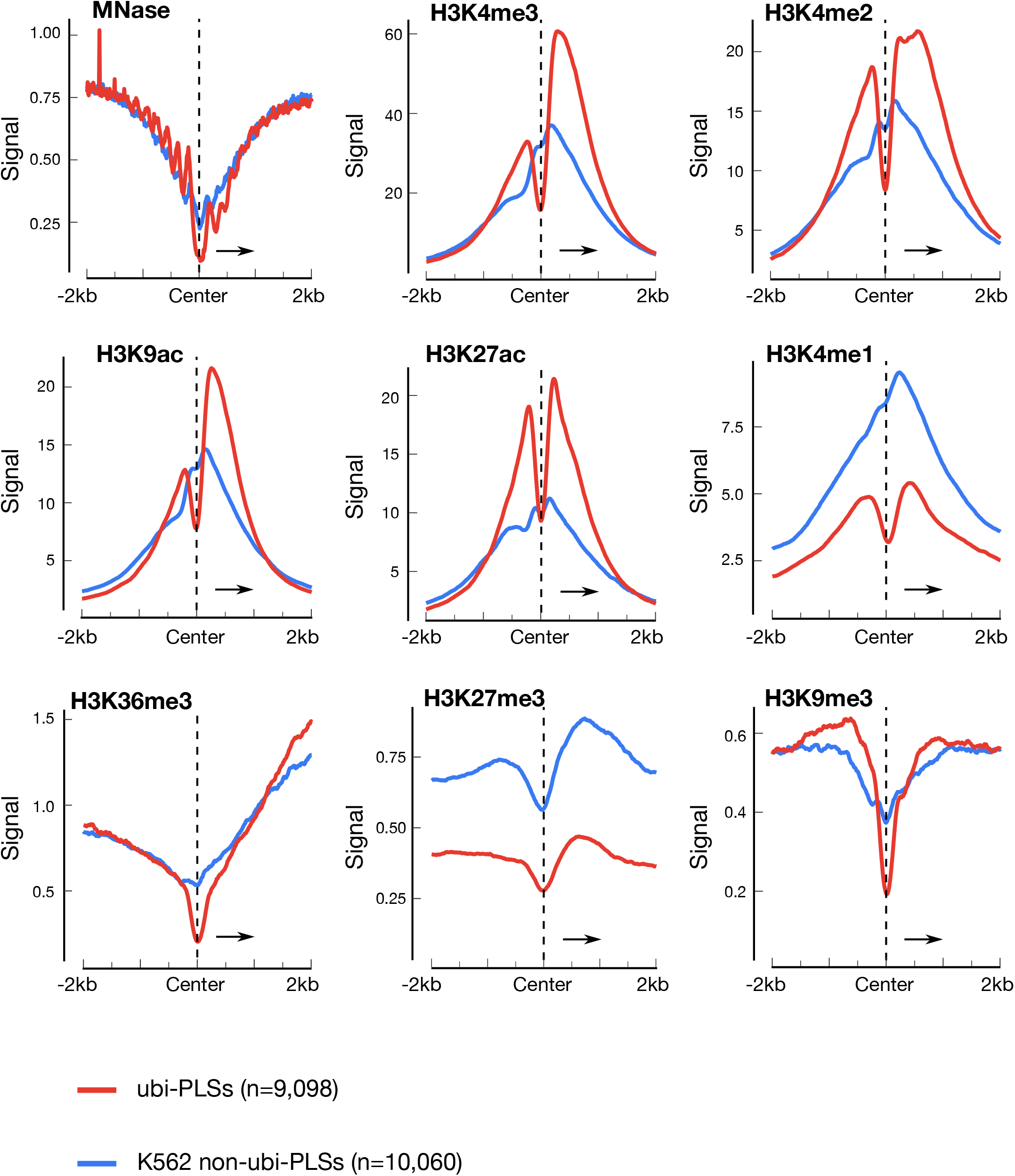
ubi-PLSs have higher signals of active histone marks and lower signals of repressive histone marks than non-ubi-PLSs. Aggregation plots depict the signal profiles of histone marks and MNase in genomic regions centered on ubi-PLSs (red) versus non-ubi-PLSs (blue) in K562 cells. Active histone marks include H3K4me3, H3K4me2, H3K27ac, H3K9ac, H3K4me1, and H3K36me3, while repressive histone marks include H3K27me3 and H3K9me3. All histone marks signals were obtained from ChIP-seq data, quantified by fold change over control. Arrows in the figure represent the transcriptional direction of the associated genes.

**Supplementary Figure 3.**
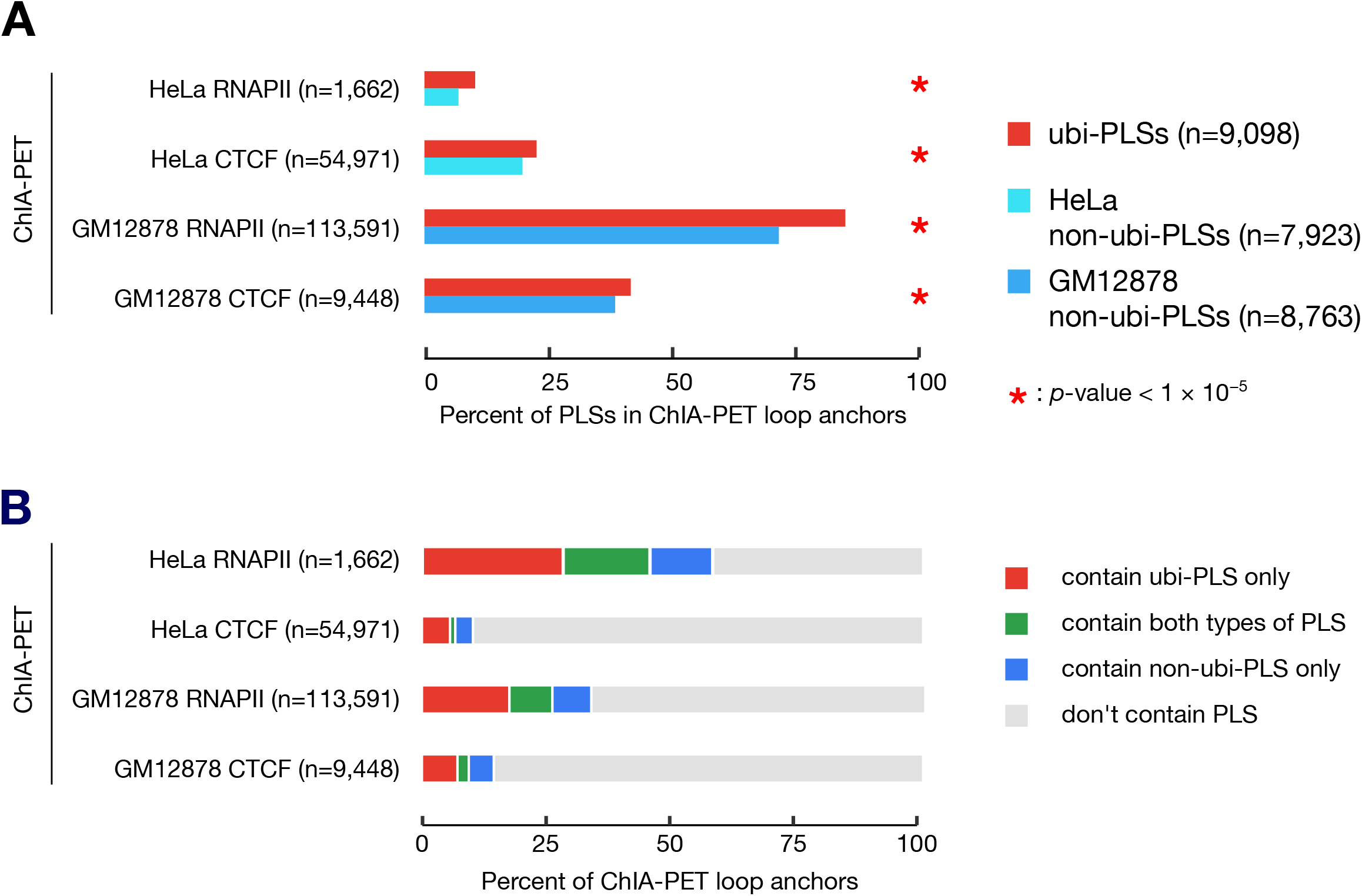
ubi-PLSs are enriched in ChIA-PET loop anchors. **A.** Higher percentages of ubi-PLSs are located in the loop anchors defined by ChIA-PET than the non-ubi-PLSs in the same cell type as the ChIA-PET data. Four ChIA-PET datasets were used: RNA Pol II (RNAPII) in HeLa cells, CTCF in HeLa cells, RNAPII in GM12878 cells, and CTCF in GM12878 cells. ubi-PLSs are shown in red, while non-ubi-PLSs are shown in different shades of blue. All *p*-values were computed with Fisher’s exact tests. **B**. A bar plot shows that higher percentages of ChIA-PET loop anchors contain ubi-PLSs than non-ubi-PLSs. For each ChIA-PET dataset, the plot shows loop anchors that only contain ubi-PLSs (red), contain both ubi-PLSs and non-ubi-PLSs (green), only contain non-ubi-PLSs (blue), or do not contain any PLSs defined in that cell type (gray).

**Supplementary Figure 4.**
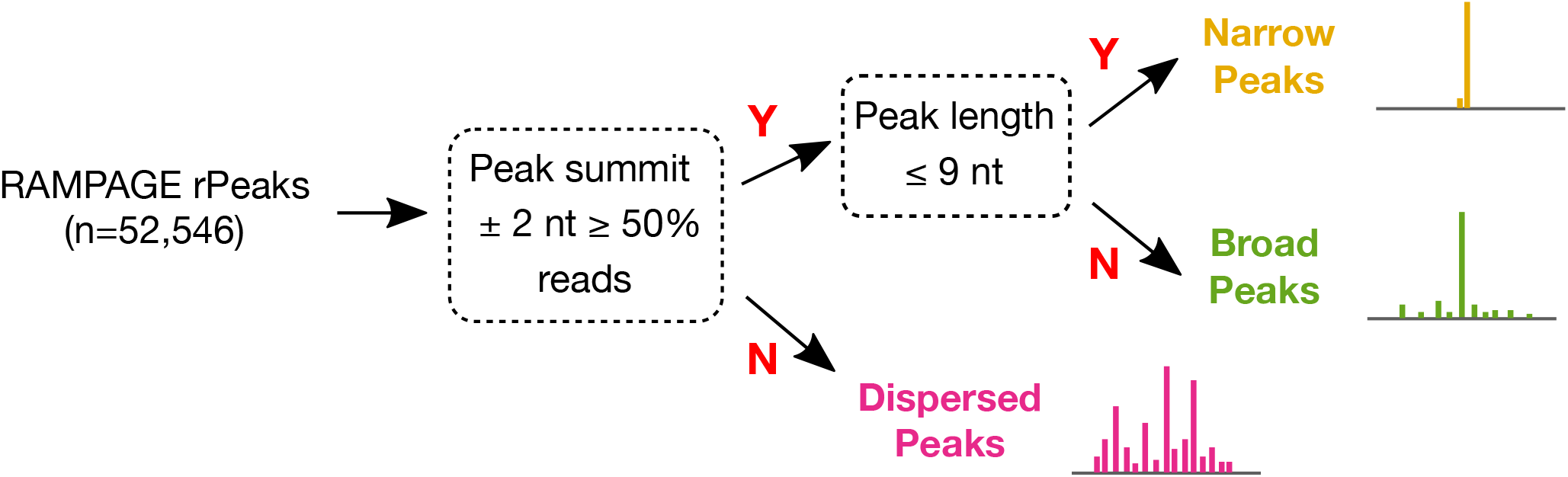
Workflow for defining promoter shapes (broad peaks, narrow peaks, and dispersed peaks).

**Supplementary Figure 5.**
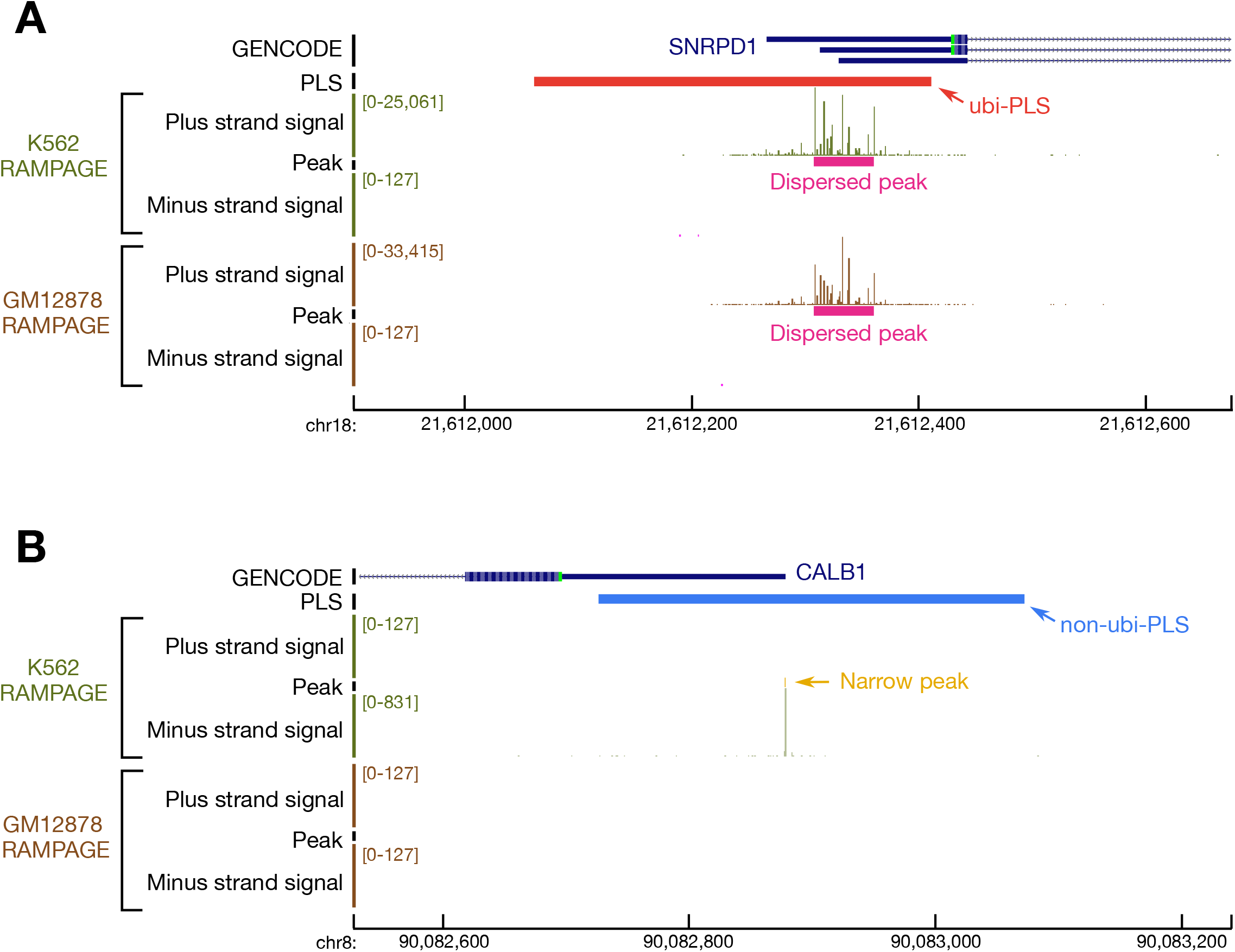
Examples of ubi-PLSs and non-ubi-PLSs with different promoter shapes. **A.** A ubi-PLS overlaps with a dispersed peak in K562 and GM12878 cells. **B.** A non-ubi-PLS overlaps with a narrow peak in K562 cells, but this TSS is not active in GM12878 cells.

**Supplementary Figure 6.**
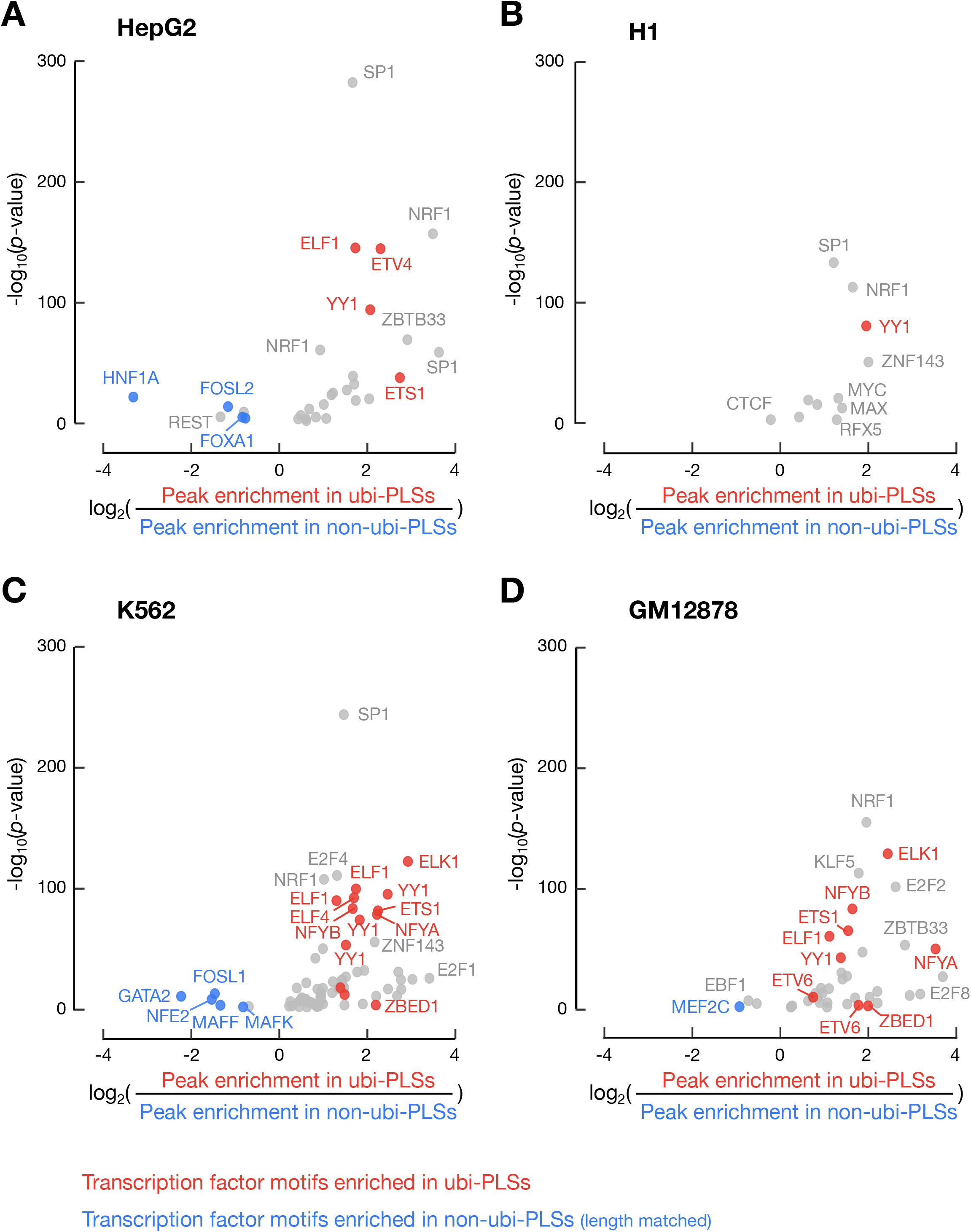
Different transcription factors bind ubi-PLSs than non-ubi-PLSs. Each dot is a transcription factor with ChIP-seq data (transcription factors with multiple ChIP-seq datasets are represented by multiple dots, e.g., SP1 in HepG2). Similar to **Figure 4B**, but the enrichment of a transcription factor is computed using their ChIP-seq peak with a motif site in HepG2 (**A**), H1 (**B**), K562 (**C**), and GM12878 (**D**) cells. As in **Figure 4B**, transcription factors whose motifs are enriched in ubi-PLSs are in red, while transcription factors whose motifs are enriched in non-ubi-PLSs are in blue. Only transcription factors with enrichment *p*-values < 0.01 are shown.

**Supplementary Figure 7.**
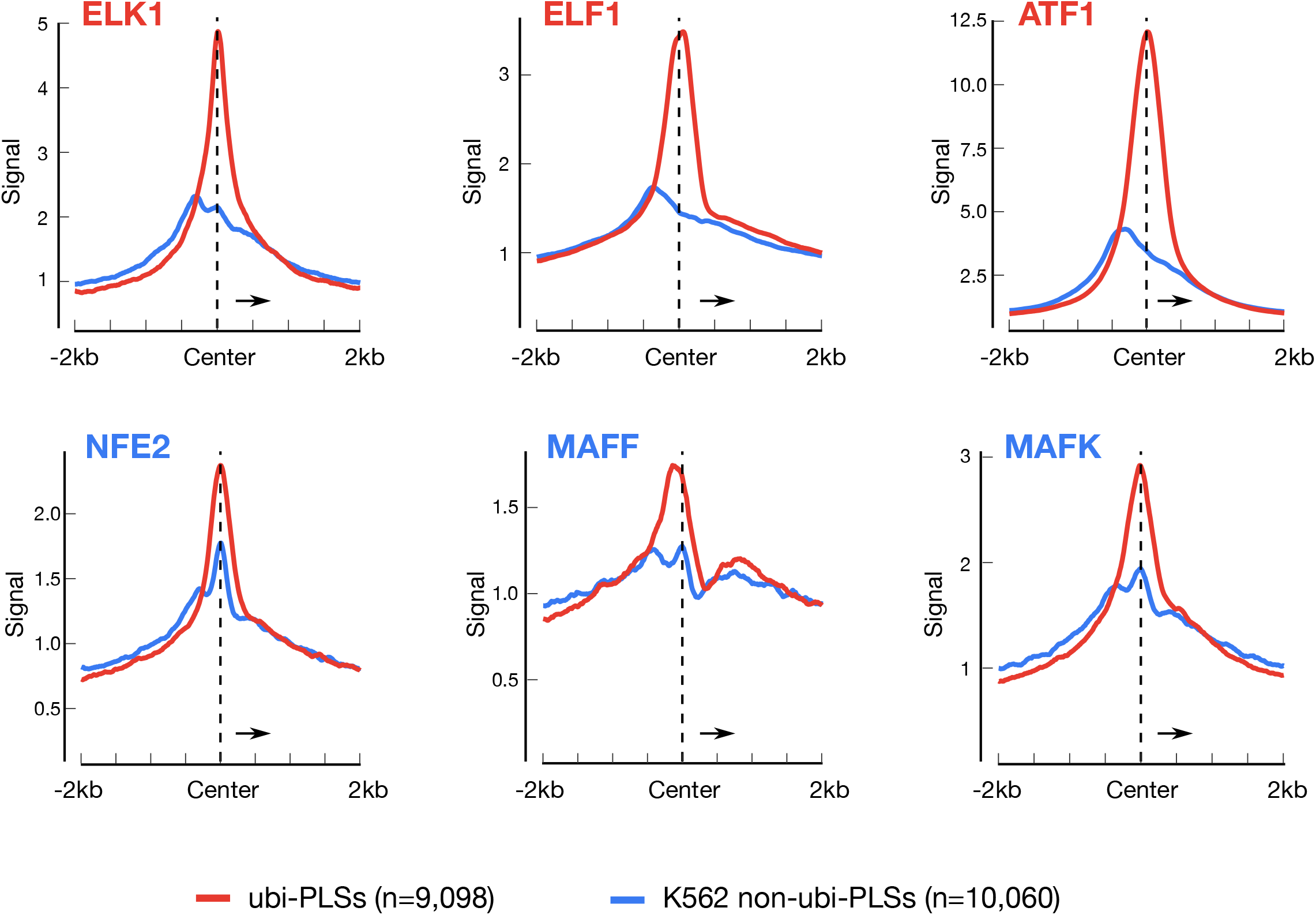
Transcription factors prefer to bind ubi-PLSs rather than non-ubi-PLSs. Aggregation plots show the ChIP-seq signal profiles for six example transcription factors in the genomic regions centered on ubi-PLSs (red) and K562 non-ubi-PLSs (blue). (top) ELK1, ELF1, and ATF1 were among the transcription factors with motifs more enriched in ubi-PLSs than in non-ubi-PLSs. (bottom) NFE2, MAFF, and MAFK were among the transcription factors with motifs more enriched in non-ubi-PLSs than in ubi-PLSs.

**Supplementary Figure 8.**
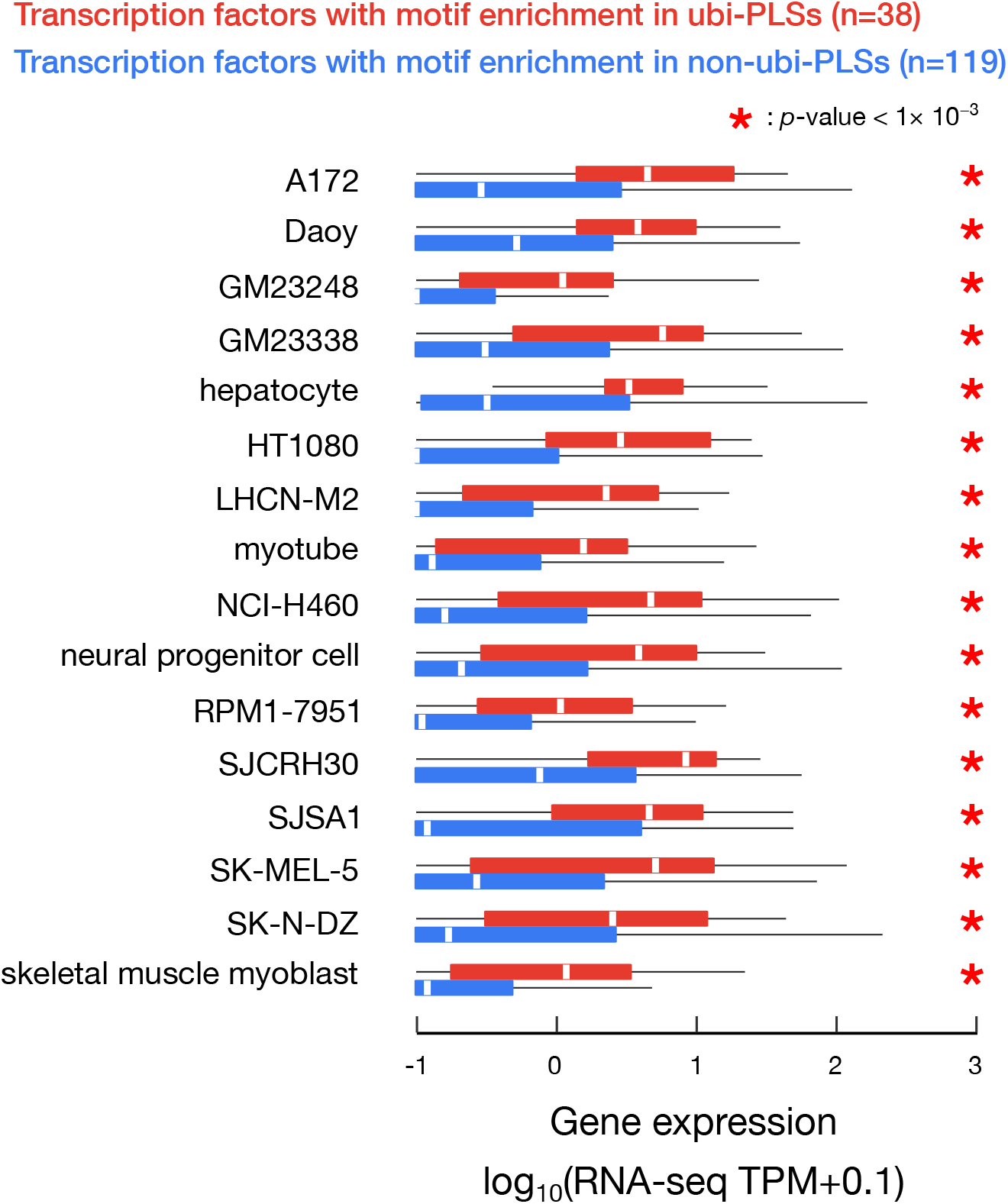
Transcription factors with motif enrichment in ubi-PLSs have higher expression levels than transcription factors with motif enrichment in non-ubi-PLSs. Boxplots show that the 38 transcription factors with motif enrichment in ubi-PLSs (red) were significantly more highly expressed than the 119 transcription factors with motif enrichment in non-ubi-PLSs (blue) in the same biosample. Expression levels were obtained from RNA-seq data, quantified in TPM (with a pseudocount of 0.1 added to each transcription factor gene), and plotted in log scale. All *p*-values were computed with Wilcoxon rank-sum tests.

## Supplementary Table Legends

**Supplementary Table 1. Accessions for ENCODE datasets used in this study.**

Spreadsheet including accessions for ENCODE human rDHS (**A**), mouse rDHS (**B**), DNase-seq (**C**), RNA-seq (**D**), RAMPAGE (**E**), histone ChIP-seq (**F**), MNase-seq (**G**), and transcription factor ChIP-seq (**H**) datasets.

**Supplementary Table 2. Lists of ubi-rDHSs as well as genes and TSSs overlapping ubi-PLSs in human (A-C) and mouse (D-F).**

Lists of ubi-rDHSs in human (**A**) and mouse (**D**), genes with TSSs overlapping human ubi-PLSs (**B**) and mouse ubi-PLSs (**E**), and TSSs overlapping human ubi-PLSs (**C**) and mouse ubi-PLSs (**F**).

**Supplementary Table 3. Genes whose TSSs overlapped ubi-PLSs are enriched in basic cellular functions.**

Gene Ontology analysis on genes whose TSSs overlapped ubi-PLSs revealed enriched Biological Process (**A**), Molecular Function (**B**), and Cellular Component (**C**). Fisher’s exact test *p*-values after FDR correction are reported.**Supplementary Table 4. Lists of transcription factors with motif enrichment in ubi-PLSs (A) and non-ubi-PLSs (B).**

## Acknowledgments

The authors thank the Weng lab for helpful feedback and discussions, and Shaimae Elhajjajy for editing the manuscript.

## Funding

This work was funded by grants from the National Institutes of Health (5U41HG007000, 5U24HG009446) to Z.W.

## Author Contributions

This project was developed and designed by K.F., J.M., and Z.W.. K.F. and X.Z. performed the analyses. K.F. and Z.W. wrote the manuscript. All authors read and approved the final version of the manuscript.

## Competing Interests

The authors declare no competing interests.

## Notes

### Competing Interest Statement

The authors have declared no competing interest.

